# A phosphorylation code coordinating transcription condensate dynamics with DNA replication

**DOI:** 10.1101/2024.05.10.593572

**Authors:** Carlos Origel Marmolejo, Celina Sanchez, Juyoung Lee, Marcel Werner, Paige Roberts, Stephan Hamperl, Joshua C. Saldivar

**Affiliations:** Cancer Early Detection Advanced Research Center, Knight Cancer Institute, Oregon Health & Science University, Portland, OR 97201; Program in Biomedical Sciences, School of Medicine, Oregon Health & Science University, Portland, OR 97201; Institute of Epigenetics and Stem Cells, Helmholtz Center Munich, Munich, Germany.; Division of Oncological Sciences, Knight Cancer Institute, Oregon Health & Science University, Portland, OR 97201

**Keywords:** Transcription condensates, membrane-less compartments, ATR, CHK1, MED1, histone locus body, genome stability, DNA damage, linker histone H1.1

## Abstract

Chromatin is organized into compartments enriched with functionally-related proteins driving non-linear biochemical activities. Some compartments, *e.g.* transcription foci, behave as liquid condensates. While the principles governing the enrichment of proteins within condensates are being elucidated, mechanisms that coordinate condensate dynamics with other nuclear processes like DNA replication have not been identified. We show that at the G1/S cell cycle transition, large transcription condensates form at histone locus bodies (HLBs) in a cyclin-dependent kinase 1 and 2 (CDK1/2)-dependent manner. As cells progress through S phase, ataxia-telangiectasia and Rad3-related (ATR) accumulates within HLBs and dissolves the associated transcription condensates. Integration of CDK1/2 and ATR signaling creates a phosphorylation code within the intrinsically-disordered region of mediator subunit 1 (MED1) coordinating condensate dynamics with DNA replication. Disruption of this code results in imbalanced histone biosynthesis, and consequently, global DNA damage. We propose the spatiotemporal dynamics of transcription condensates are actively controlled via phosphorylation and essential for viability of proliferating cells.

## INTRODUCTION

The nucleus exhibits remarkable spatial organization through compartmentalization of chromatin regions where dynamic processes, such as transcription, replication, and DNA repair are active. These compartments lack membranes, yet are highly-enriched with proteins that are de-mixed from the nucleoplasm to perform biochemical functions. In the case of transcription, compartmentalization can selectively raise the local concentration of functionally-related proteins and exclude opposing factors creating non-linear gene regulatory behaviors[1].

The initiation of transcription occurs in compartments commonly referred to as transcription condensates[2, 3]. These compartments exhibit liquid-like properties and are thought to influence the 3-dimensional architecture of chromatin bringing enhancers and super-enhancers in close proximity to promoters to drive gene expression[4, 5]. Transcription condensates form as a result of weak, multivalent protein-protein and protein-RNA interactions[1]. Indeed, many transcription factors and co-activators contain intrinsically-disordered regions (IDRs) that can form multivalent interactions with other IDRs and RNA[6].

How these dynamic compartments are controlled with regards to their formation, equilibrium, and dissolution is poorly understood. Recent findings suggest the pattern of alternating blocks of negatively and positively charged amino acids within protein IDRs can specify the selective partitioning of transcriptional regulators[7]. Moreover, cell cycle-specific hyperphosphorylation of Ki-67 and NPM1 changes the patterns of alternating charge blocks in these proteins and alters their propensity for compartmentalization[8]. In addition, the kinase activity of DRK3 dissolves several types of membrane-less compartments in mitosis[9]. RNA polymerase II (RNAPII) is itself phosphorylated along its intrinsically-disordered C-terminal domain (CTD), and changes in the phosphorylated residues trigger a switch from transcription to splicing-specific condensates[5, 10]. These examples suggest phosphorylation may be a widespread mechanism to regulate condensate formation and dissolution.

In proliferating cells, regulation of transcription condensate dynamics is particularly important as cells need to coordinate their spatial organization with DNA replication to maintain replication fidelity, transcriptional states, and genome integrity[11]. In S phase, there are several kinases that are highly active and could potentially regulate condensates via phosphorylation of transcriptional regulators. These include the cyclin-dependent kinases, CDK2 and CDK1, and the S phase checkpoint kinases, ATR and CHK1, which are known regulators of replication initiation and transcription [12–17]. Here, we uncover a phosphorylation code present in the IDR of mediator subunit 1 (MED1), governed largely by the integrated activities of ATR, CHK1, CDK1 and CDK2, that controls transcription condensate dynamics and coordinates them with DNA replication to ensure genome stability and cell viability.

## RESULTS

### Large transcription condensates form at the G1/S transition and are dissolved by ATR signaling during S phase progression

To investigate the regulation of transcription condensates across S phase, we imaged MED1, a subunit of the mediator complex that plays a primary role in condensate formation, by indirect immunofluorescence confocal microscopy in the non-malignant epithelial cell lines, MCF10A and RPE-1 hTERT. We labeled the S phase population using a 15-minute pulse with EdU just prior to fixation. Consistent with previous reports[2, 18–20], MED1 formed numerous foci throughout the nucleus of interphase cells. Notably, a subset of cells, 10-20%, contained 2-5 large MED1 foci that were about a micron in diameter (Figures 1A and S1A and B). These large foci overlapped with RNAPII and BRD4 foci (Figure 1B), consistent with their being transcription condensates. Intriguingly, the large MED1 foci were only present in EdU-positive cells (Figure 1A), suggesting large transcription condensates are tightly-coupled with S phase.

**Figure 1.**
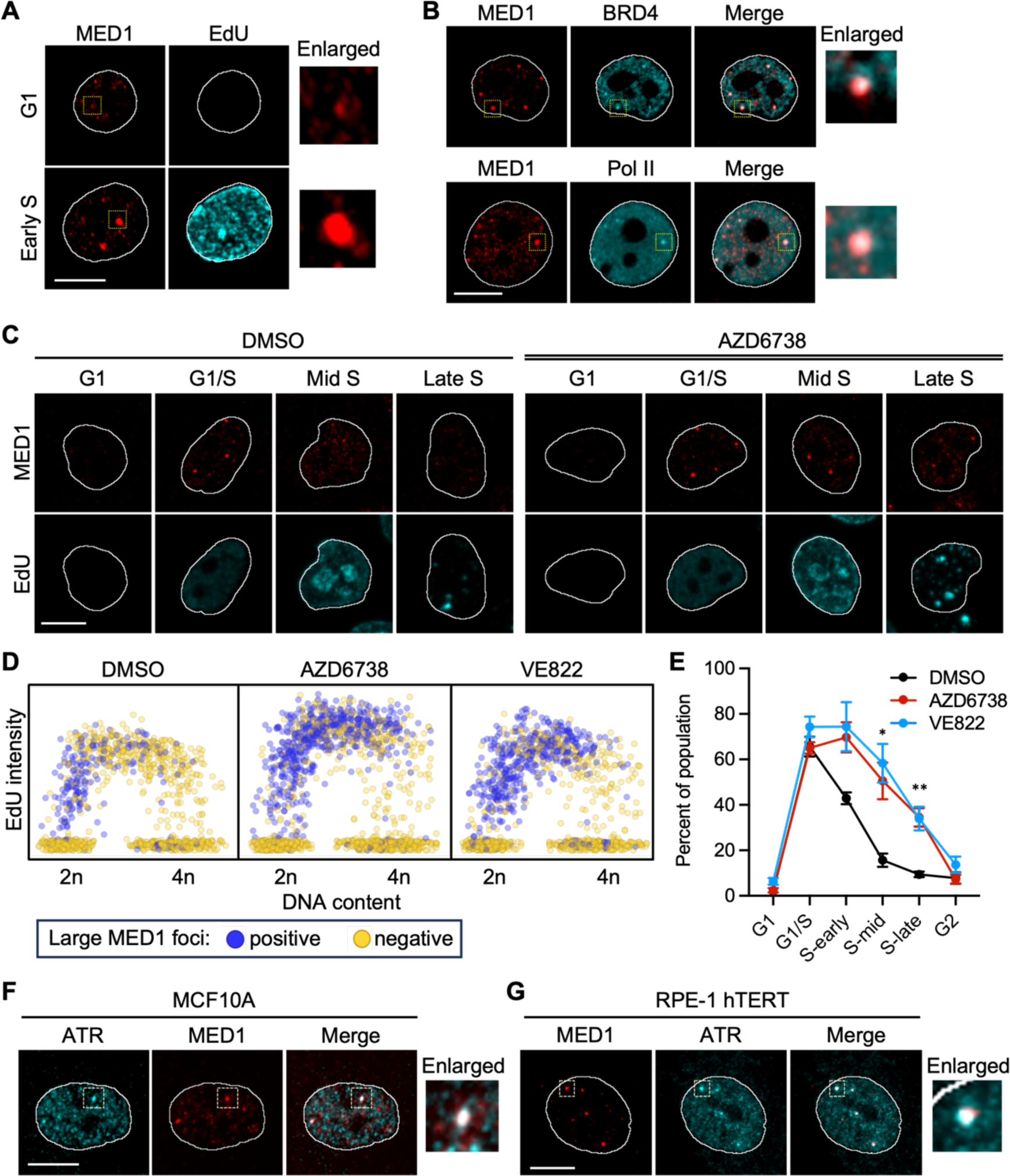
Large transcription condensates appear at the G1/S transition and are dissolved during S phase by ATR kinase activity. (A) Representative images of MED1 and EdU in MCF10A cells. Yellow boxes mark regions of enlarged condensates. Scale bar, 10 μm. (B) Representative images of MED1, BRD4, and Pol II (RNAPII) in MCF10A cells. Yellow boxes mark regions of enlarged condensates. Scale bar, 10 μm. (C) Representative images of MED1 and EdU in MCF10A cells at various points in the cell cycle. Cell cycle phase was determined by QIBC as described in Supplemental Fig 1D. Cells were treated with DMSO (mock) or 5 μM AZD6738 (ATR inhibitor) for 1 h. Scale bar, 10 μm. (D) Scatter plot of DAPI integrated intensity (linear scale) and EdU mean intensity (log_2_ scale) in MCF10A cells treated with DMSO (mock), 5 μM AZD6738 (ATR inhibitor), or 5 μM VE822 (ATR inhibitor) for 1 h. 2n and 4n denote the DNA content. Cells positive for large MED1 foci (diameter > 0.7 μm) were colored purple, and negative cells were yellow. (E) Line plots showing the percent of cells with large MED1 foci for each cell cycle phase. Cells were treated as in (D). Points and error bars are the mean and standard error, respectively, of 3 independent experiments. (* p < 0.05, ** p < 0.01). (F) Representative images of MED1 and ATR in MCF10A cells. Yellow boxes mark regions of enlarged condensates. Scale bar, 10 μm. (G) Representative images of MED1 and ATR in RPE-1 hTERT cells. Yellow boxes mark regions of enlarged condensates. Scale bar, 10 μm.

Given that ATR is linked to regulation of transcription in S phase[13], we investigated the impact of ATR inhibition on the large transcription condensates. Within 1 hour of ATR inhibition using two structurally-distinct inhibitors, AZD6738 and VE822, we observed an approximate 2-fold increase in the percent of cells that contained the large MED1 foci compared to the DMSO vehicle control (Figures S1B and S1C). To more quantitatively measure the MED1 foci across the cell cycle, we developed an automated image analysis pipeline that integrates focus detection and quantification with quantitative image-based cytometry (QIBC)[21]. QIBC allows us to identify the G1, G1/S, early S, mid-S, late-S, and G2 populations (Figure S1D), and the focus detection allows us to measure the number of MED1 foci and foci intensities for each cell across the different cell cycle phases (Figure S1E). Strikingly, the large foci appeared at the G1/S transition, and then gradually disappeared as cells progressed into mid- and late-S phase (Figures 1C-E). Inhibition of ATR for 1 h with either AZD6738 or VE822 caused the MED1 foci to persist into late S phase. Moreover, co-staining with siRNA-validated antibodies (Figure S1F) revealed a prominent co-localization of large ATR foci with the large MED1 foci in MCF10A cells (Figure 1F), RPE-1 hTERT cells (Figure 1G), and in HBEC-3KT cells (Figure S1G), suggesting the ATR-MED1 compartment is common across different cell types. Of note, the smaller ATR and MED1 foci did not co-localize. In addition, a proximity ligase assay using ATR and RNAPII antibodies revealed that ATR and RNAPII are in close proximity in a transcription-dependent manner (Figure S1H). Collectively, these results suggest ATR accumulates within large transcription condensates that form at the G1/S transition and dissolves them during S phase progression via phosphorylation.

### S phase transcription condensates form at the histone locus body

Next, we asked whether ATR enrichment within the S phase transcription condensates was due to DNA damage. Notably, the large MED1 foci did not colocalize with yH2AX or RPA32 foci (Figure S2A), evidence that ATR is not recruited to the large condensates by DNA damage or RPA. Given the size of the ATR-MED1 foci and our observation that there were 2-5 per cell, we reasoned the large condensates may instead form at nuclear bodies. The MED1 foci occasionally formed adjacent to the Cajal body marker, coilin, a spatial pattern that resembles the histone locus body (HLB)[22] (Figure S2B). HLBs are nuclear compartments that form around the replication-dependent histone gene clusters and function to couple histone gene expression and RNA processing to S phase[23–25]. Indeed, co-staining MED1 or ATR with NPAT, a scaffold protein that marks HLBs, revealed that the large ATR-MED1 foci form at HLBs (Figures S2C and S2D).

Using our focus detection and QIBC platform, we found that MED1 levels abruptly rise within HLBs at the G1/S transition and then gradually decrease in an ATR-dependent manner during S phase progression (Figures 2A-C, S2E and S2F). By contrast, ATR levels in HLBs were lowest at the G1/S transition but then gradually increased from mid-S phase to the G2 phase (Figures 2D and E). We speculate that the rise in ATR abundance at HLBs, as opposed to an increase in ATR kinase activity, leads to dissolution of the transcription condensate from the nuclear body. To our knowledge, this is the first time ATR has been associated with the HLB and implicate ATR and MED1 as dynamic components of this nuclear compartment.

**Figure 2.**
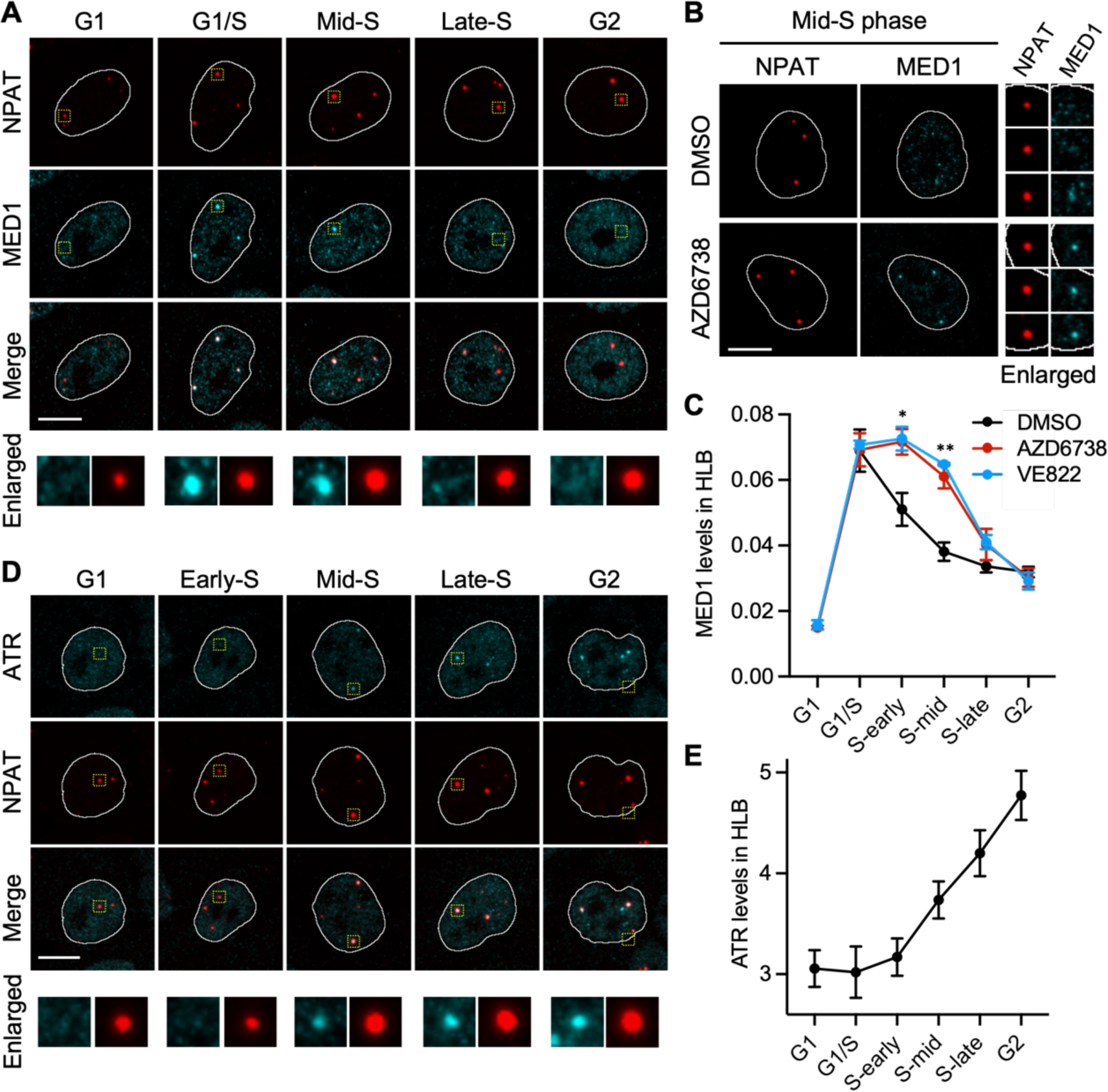
The S-phase transcription condensates form at the histone locus bodies. (A) Representative images of MED1 and NPAT in MCF10A cells at various points in the cell cycle. Yellow boxes mark regions of enlarged condensates. Scale bar, 10 μm. (B) Representative images of MED1 and NPAT in mid-S phase MCF10A cells. Cells were treated with DMSO (mock) or 5 μM AZD6738 (ATR inhibitor) for 1 hour. Each NPAT-marked HLB focus was enlarged. Scale bar, 10 μm. (C) Line plots showing the levels of MED1 in NPAT-marked HLBs measured by immunofluorescence. Points and error bars are the mean and standard error, respectively, of 3 independent experiments. (* p < 0.05, ** p < 0.01). (D) Representative images of ATR and NPAT in MCF10A cells at various points in the cell cycle. Yellow boxes mark regions of enlarged condensates. Scale bar, 10 μm. (E) Line plots showing the levels of ATR in NPAT-marked HLBs measured by immunofluorescence. Points and error bars are the mean and standard error, respectively, of 3 independent experiments.

### ATR signaling increases the liquid-like property of the S phase transcription condensates

Transcription condensates are thought to form through multivalent, weak protein-protein and protein-RNA interactions[1]. Condensates exist on a continuum across a liquid state, a gel-like state, and a solid state, depending on the electrostatic properties of the multivalent interactions (weaker affinities in the liquid state and stronger affinities in the solid state)[1]. To investigate the electrostatic properties of the large MED1 foci, we used the aliphatic alcohol 1,6-hexanediol (1,6-HXD), which dissolves condensates by disrupting their weak multivalent interactions[26]. Consistent with previous reports[2, 26], MED1 foci were disrupted within 1 minute by increasing concentrations of 1,6-HXD (Figures 3A and 3B). Examination of the effect of 1,6-HXD on MED1 foci across the cell cycle revealed a striking impact on MED1 foci at HLBs in S phase (Figure 3C). Notably, the G1/S foci, and to a lesser degree the early S phase foci, were more resistant to disruption by 1,6-HXD compared to mid- and late-S phase foci (Figure 3D). Furthermore, in mid-S phase when the large MED1 foci dissolve in an ATR-dependent manner, we observed specifically these large foci (0.4-0.6 μm^2^ area) to be the most sensitive to 1,6-HXD treatment (Figure 3E). These findings suggest the affinities of MED1 with other components within transcription condensates at HLBs are highest when they first appear at the start of S phase and then decrease as cell progress through S phase, resulting in a transition from more solid-like compartments to more liquid-like before they dissolve in mid-S phase.

**Figure 3.**
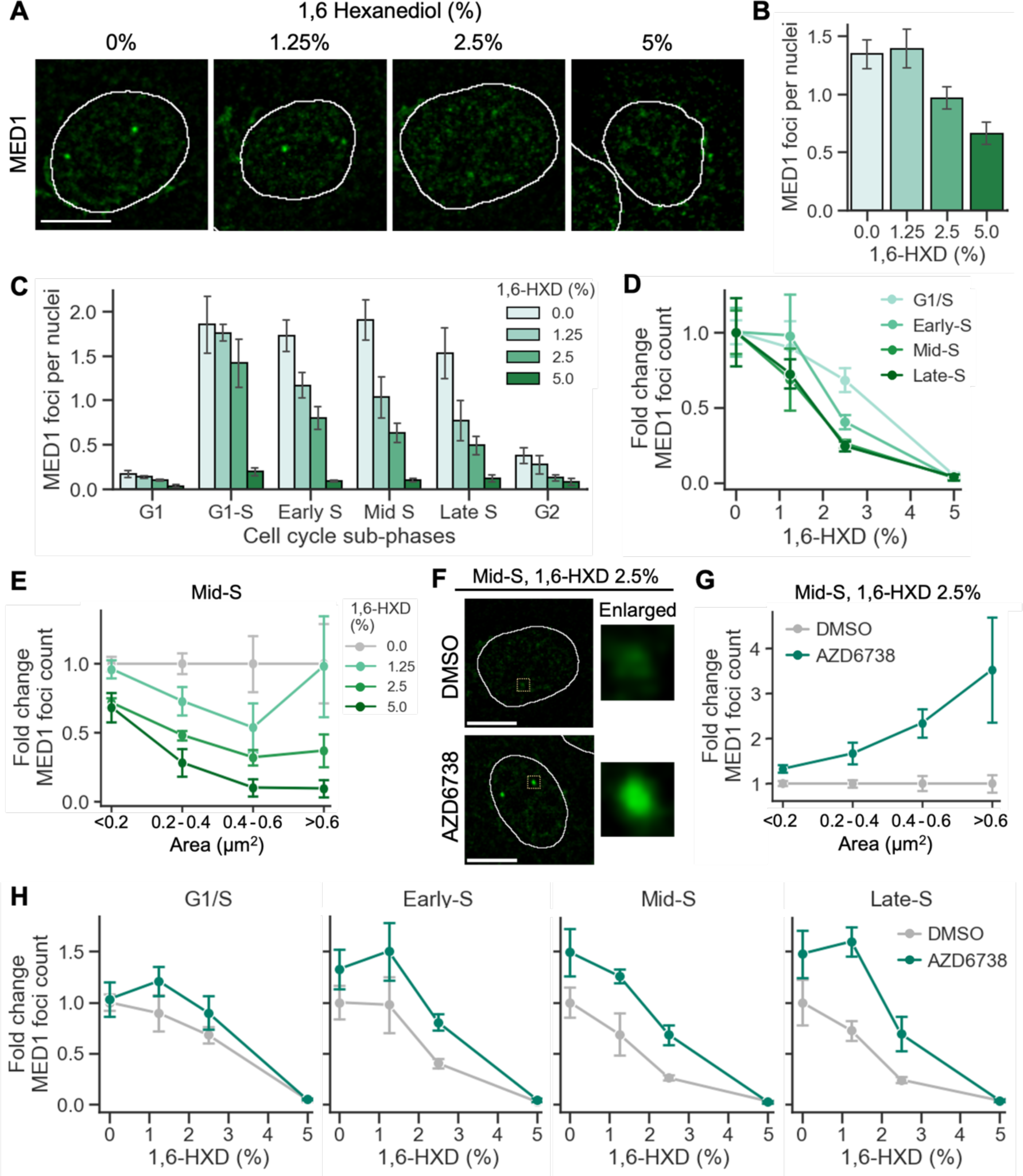
ATR increases the liquid-like property of S-phase transcription condensates. (A) Representative images of MED1 in mid-S phase MCF10A cells. Cells were treated with 0, 1.25, 2.5, or 5 % 1,6 hexanediol for 1 min. Scale bar, 10 μm. (B) Bar plot showing the quantification of MED1 foci (diameter > 0.7 μm) in cells treated with varying concentrations (0%, 1.25%, 2.5%, and 5%) of 1,6-hexanediol (1,6-HXD) for 1 min. Bars and error bars are the mean and standard error, respectively, of 3 independent experiments. (C) Bar plot showing the quantification of MED1 foci (diameter > 0.7 μm) in cells treated with varying concentrations (0%, 1.25%, 2.5%, and 5%) of 1,6-HXD for 1 min at different stages of the cell cycle. The color of each bar corresponds to the concentration of 1,6-HXD used. Bars and error bars are the mean and standard error, respectively, of 3 independent experiments. (D) Line plot showing the fold change in the number of MED1 foci following a 1 min treatment with 1.25, 2.5, or 5 % 1,6-HXD compared to 0% 1,6-HXD. The fold change is calculated across G1/S, early-S, mid-S, and late-S phase. Each line represents a different cell cycle phase. Cells were treated as in (C). Points and error bars are the mean and standard error, respectively, of 3 independent experiments. (E) Line plot illustrating the fold change in the number of MED1 foci following a 1 min treatment with 1.25, 2.5, and 5 % 1,6-HXD compared to 0% 1,6-HXD. The fold change is calculated across four distinct groups of MED1 foci categorized by area: <0.2, 0.2 to 0.4, 0.4 to 0.6, and >0.6 µm². Each line represents a different 1,6-HXD concentration. Points and error bars are the mean and standard error, respectively, of 3 independent experiments. (F) Representative images of MED1 in mid-S phase cells treated with 2.5 % 1,6-HXD for 1 min. Yellow boxes mark regions of enlarged condensates. Scale bar, 10 μm. (G) Line plot showing the fold change in the number of MED1 foci in mid-S phase cells after a 1 h treatment with 5 µM AZD6738 compared to DMSO. Cells were also treated with 2.5 % 1,6-HXD. The fold change is calculated across four distinct groups of MED1 foci categorized by area: <0.2, 0.2 to 0.4, 0.4 to 0.6, and >0.6 µm². Each line represents a different treatment condition (DMSO or AZD6738). Points and error bars are the mean and standard error, respectively, of 3 independent experiments. (H) Line plot showing the fold change in the number of MED1 foci in cells treated with DMSO or 5 μM AZD6738 for 1 h, followed by a 1 min treatment with 1.25, 2.5, and 5 % 1,6-HXD compared to 0% 1,6-HXD. The fold change is calculated across G1/S, early-S, mid-S, and late-S phase. Each line represents a different treatment condition (DMSO or AZD6738). Points and error bars are the mean and standard error, respectively, of 3 independent experiments.

Next, we tested the effect of 1,6-HXD on NPAT and ATR foci. Consistent with NPAT acting as a scaffold for the HLB, the NPAT foci were largely unaffected by treatment with 1,6-HXD except for at the highest concentration (Figures S3A-S3D). Curiously, 1,6-HXD increased the number of ATR foci throughout the cell cycle irrespective of focus size (Figures S3E-H). Thus, while NPAT and ATR are key components of the HLB, their enrichment in the compartment is likely not driven by multivalent interactions between IDRs like that of MED1. Accordingly, the HLB is a compartment that contains both structural components (NPAT), dynamic droplet-like components (MED1), and dynamic components with unknown features (ATR).

Given that mid-S phase condensates dissolve from HLBs in an ATR-dependent manner, we asked whether ATR influenced the electrostatic properties of MED1. Strikingly, the large MED1 foci in mid-S phase became resistant to 1,6-HXD treatment after 1 hour inhibition of ATR (Figures 3F and 3G). Resistance to 1,6-HXD became more prominent upon ATR inhibition in cells that were further into S phase, with the G1/S population largely unaffected by ATR inhibition and the late-S phase population most strongly affected (Figure 3H). We conclude that ATR kinase activity lowers the affinity among the multivalent interactions within the HLB transcription condensates as cells progress through S phase making them more liquid-like and ultimately dissolving them.

### Cell cycle checkpoint signaling controls S phase transcription condensate dynamics

To investigate the mechanism through which ATR controls MED1 condensate dynamics across S phase, we asked whether CHK1, a downstream effector kinase of ATR that regulates the S phase cell cycle checkpoint[12], also controlled MED1 dynamics. Using two different CHK1 inhibitors (LY2603618 and ChIR-124) we found that, like ATR, CHK1 inhibition resulted in the persistence of MED1 foci at HLBs into late S phase (Figures 4A and 4B), suggesting ATR controls transcription condensate dynamics through its downstream effector CHK1.

**Figure 4.**
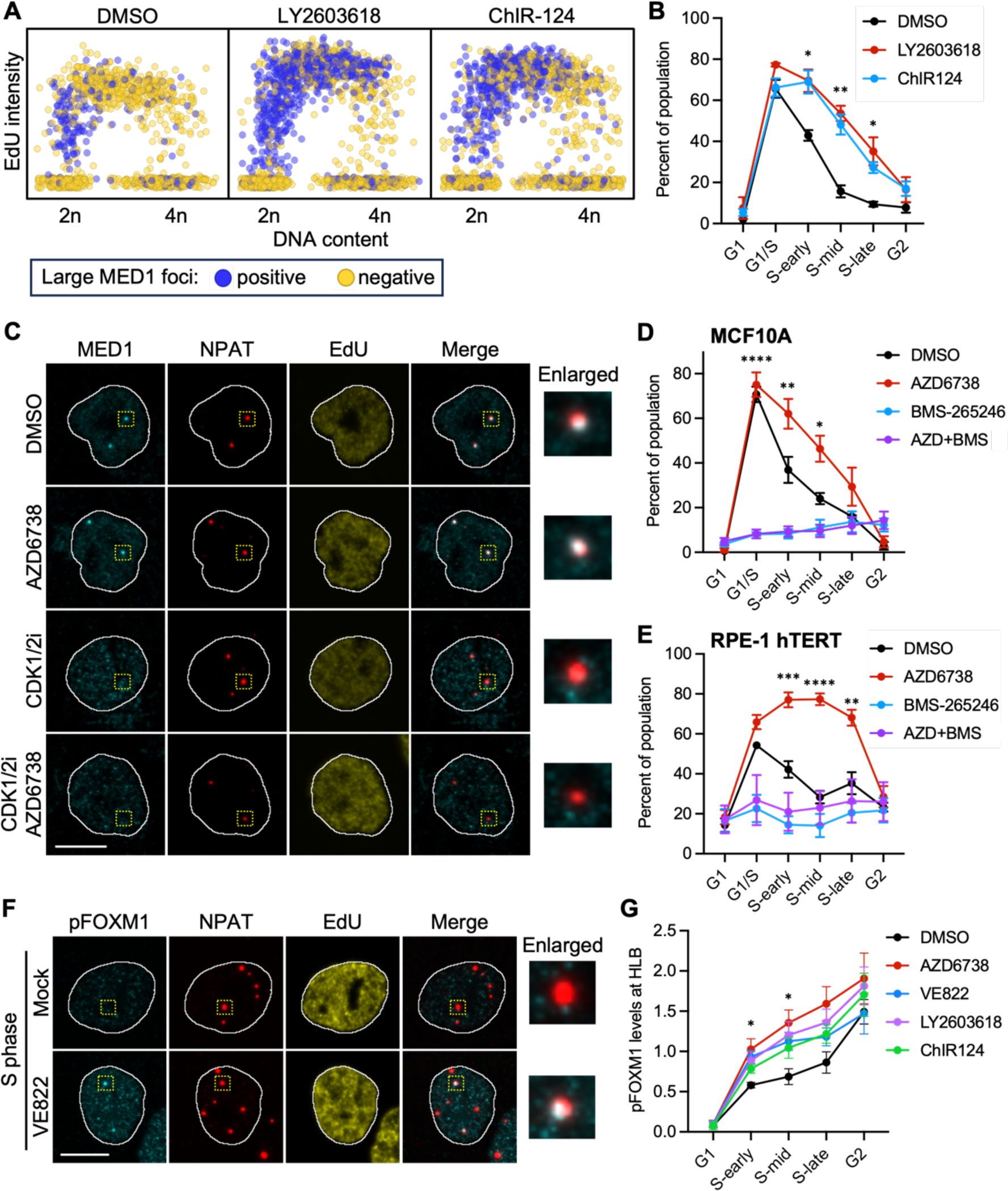
S-phase checkpoint signaling controls transcription condensate dynamics. (A) Scatter plot of DAPI integrated intensity (linear scale) and EdU mean intensity (log_2_ scale) in MCF10A cells treated with DMSO (mock), 2 μM LY2603618 (CHK1 inhibitor), or 250 nM ChIR-124 (CHK1 inhibitor) for 1 h. 2n and 4n denote the DNA content. Cells positive for large MED1 foci (diameter > 0.7 μm) were colored purple, and negative cells were yellow. (B) Line plots showing the percent of cells with large MED1 foci for each cell cycle phase. Cells were treated as in (A). Points and error bars are the mean and standard error, respectively, of 3 independent experiments. (* p < 0.05, ** p < 0.01). (C) Representative images of MED1, NPAT, and EdU in early S-phase MCF10A cells treated with DMSO (mock), 5 μM AZD6738 (ATR inhibitor), 5 μM BMS-265246 (CDK1/2 inhibitor), or both the ATR inhibitor and the CDK1/2 inhibitor for 1 h. Yellow boxes mark regions of enlarged condensates. Scale bar, 10 μm. (D,E) Line plots showing the percent of MCF10A cells (D) and RPE1-hTERT cells (E) with large MED1 foci for each cell cycle phase. Cells were treated as in (C). Points and error bars are the mean and standard error, respectively, of 3 independent experiments. (* p < 0.05, ** p < 0.01, *** p < 0.001, **** p < 0.0001). (F) Representative images of pFOXM1 (T600), NPAT, and EdU in MCF10A cells treated with DMSO (mock) or 5 μM VE822 (ATR inhibitor) for 1 hour. Yellow boxes mark regions of enlarged condensates. Scale bar, 10 μm. (G) Line plots showing the levels of pFOXM1 (T600) in NPAT-marked HLBs measured by immunofluorescence. Cells were treated with DMSO (mock), 5 μM AZD6738 (ATR inhibitor), 5 μM VE822 (ATR inhibitor), 2 μM LY2603618 (CHK1 inhibitor), or 250 nM ChIR-124 (CHK1 inhibitor) for 1 hour. Points and error bars are the mean and standard error, respectively, of 3 independent experiments. (* p < 0.05).

ATR-CHK1 signaling during S phase limits the kinase activities of CDK1 and CDK2 (CDK1/2)[12, 17, 27]. Accordingly, the persistence of MED1 foci into late S phase may be caused by hyperactive CDK1 and/or CDK2. Individual inhibition of either CDK1 or CDK2, with RO-3306 or NU6140 respectively, had little impact on MED1 foci (Figures S4A and S4B). By contrast combined inhibition of CDK1/2, with BMS-265246, resulted in a near complete loss of large MED1 condensates in S phase, even in cells with ATR inhibition (Figures 4C-E). These data suggest that CDK1/2 signals the formation of the large transcription condensates at the G1/S transition, and then ATR-CHK1 subsequently signals their dissolution.

We were unable to detect CHK1 foci that colocalized with MED1 or HLBs. However, unlike ATR, CHK1 is thought to diffuse away from chromatin bound by ATR[28]. Thus, CHK1 may not accumulate within HLBs at levels detectable by our imaging platform. Nevertheless, the CHK1 and CDK1/2 inhibitor data implicate HLBs as compartments where the ATR-CHK1 pathway suppresses CDK1/2 activity. In support of this, we observed phosphorylation of FOXM1 at T600 (pFOXM1), a CDK1-dependent event, to be focally-enriched at HLBs (Figures 4F and 4G). FOXM1 is normally phosphorylated in the G2 phase but is prematurely phosphorylated by CDK1 in S phase when the ATR-enforced S/G2 checkpoint is inhibited[13, 29, 30], and we observed this premature phosphorylation to also occur within HLBs (Figures 4F and 4G). This suggests ATR-CHK1 signaling is suppressing CDK1 within this compartment. Based on the collective results, we conclude that integration of ATR-CHK1 and CDK1/2 kinase signaling controls the formation and dissolution of the HLB transcription condensates during S phase.

### A MED1^IDR^ phosphorylation code couples transcription condensate dynamics to cell cycle progression

MED1 transcription condensates form through a process dependent on its large IDR that makes up the C-terminal half of the protein[2]. It is thought the unstructured IDR can form multivalent interactions with other IDRs and RNAs to drive liquid-liquid phase separation leading to transcription condensate formation[6]. To confirm the role of the MED1 IDR (MED1^IDR^) in condensate formation, we cloned both MED1^IDR^ and the fluorescent mNeonGreen protein to create a hybrid MED1^IDR^-mNeonGreen under the control of the doxycycline-inducible, TetON promoter. We generated stable MCF10A cell lines with either the inducible MED1^IDR^-mNeongreen (WT-IDR-NG) or an inducible mNeonGreen control lacking MED1^IDR^ (control-NG). Two days after induction, live-imaging of the fluorescent mNeonGreen reporter revealed WT-IDR-NG to form foci in cells when expressed above a certain threshold (Figures S5A and S5B). Control-NG did not form foci, confirming MED^IDR^ as the trigger for focus formation.

Next, we hypothesized ATR-CHK1 and CDK1/2 signaling control S phase MED1 condensates through a mechanism involving MED1^IDR^. Notably, MED1^IDR^ is highly phosphorylated, with many phosphorylation sites residing within CDK1/2, ATR, and CHK1 consensus phosphorylation motifs[31, 32]. As phosphorylation changes the charge of proteins and influences electrostatic forces between molecules[33], we reasoned multi-site phosphorylation may control condensate formation and/or dissolution. A phospho-proteomic study looking at quantitative changes in protein phosphorylation across the cell cycle revealed striking changes in MED1^IDR^ phosphorylation at the G1/S transition, in late S phase, and in the G2 phase[32]. Many of the phosphorylation changes were clustered, and considering them as phosphorylation domains (Figure 5A), we identified a distinct pattern across the cell cycle (Figure 5B and Table S1). For example, in the G1 phase, only phosphorylation domain 4 (PD 4) was hyper-phosphorylated. Upon the G1/S transition, only domain 2 (PD 2) was hyper-phosphorylated. By late S phase, domains 2 and 4 were both hyper-phosphorylated (PD 2,4). Finally, in G2 phase cells, domains 4 and 5 were hyper-phosphorylated (PD 4,5) (Figure 5B). When compared with the MED1 condensate dynamics (Figure 1E), the pattern of phosphorylation suggested the possibility of a phosphorylation code controlling the formation and dissolution of MED1 condensates across S phase.

**Figure 5.**
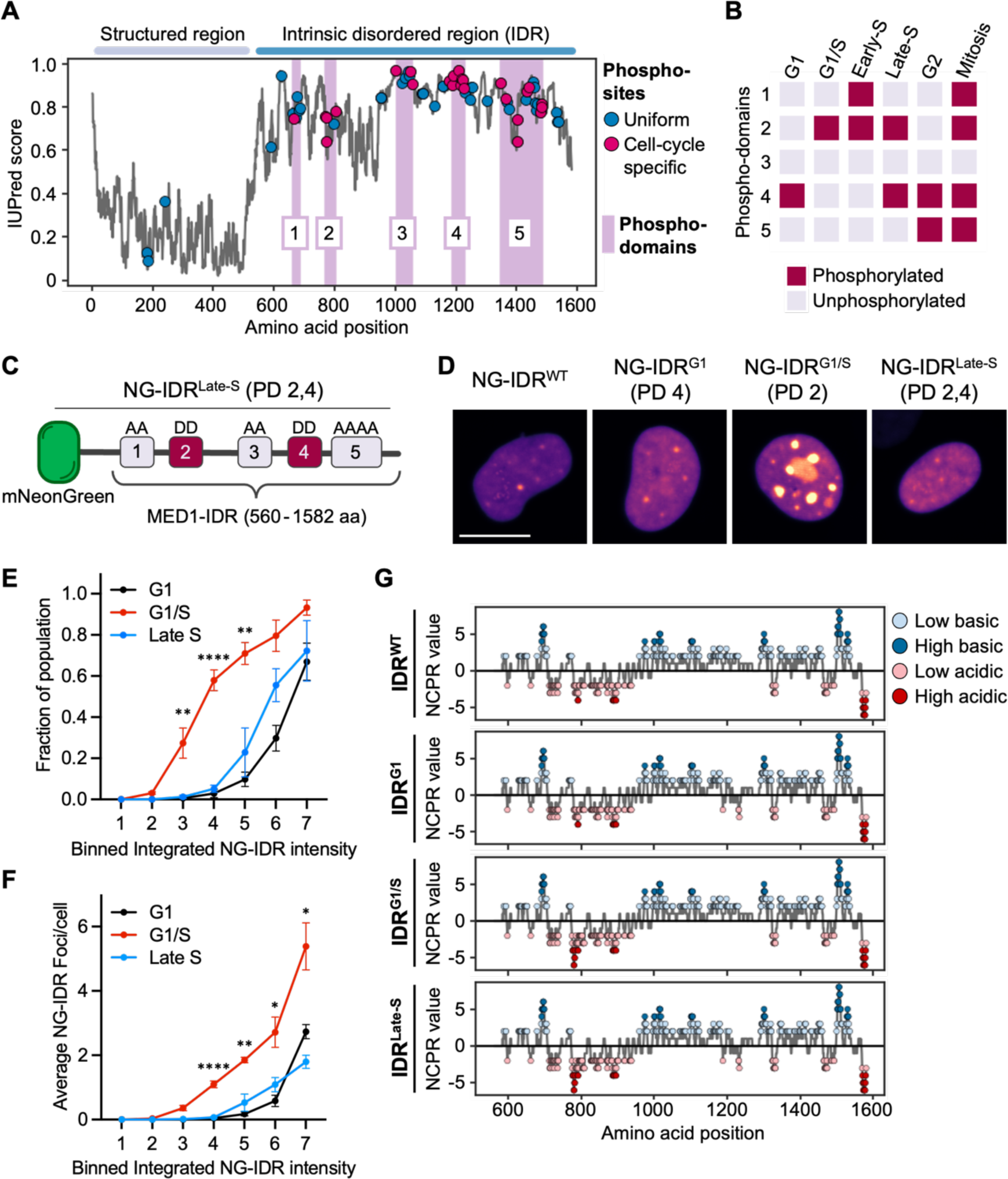
A MED1^IDR^ phosphorylation code couples transcription condensate dynamics to cell cycle progression. (A) Line plot illustrating the intrinsic unstructured protein prediction score for each residue of the MED1 sequence with the observed phosphorylation sites in the protein. Defined regions of the protein, structured and intrinsic disordered regions, are indicated on top of the plot. Color coded phosphorylation sites indicate if the phosphorylation event was detected uniformly throughout the cell cycle (uniform) or they were specific to a cell cycle phase (cell-cycle specific). Purple rectangles indicate the defined phospho-domains (1, 2, 3, 4, and 5). (B) Heatmap showing the phosphorylation status of the phospho-domains (1, 2, 3, 4, and 5) across the cell cycle (G1, G1/S, Early-S, Late-S, G2, Mitosis). (C) Schematic representation of the fluorescent reporter protein, mNeonGreen (NG), fused to the MED1-IDR (560 - 1582 aa) with amino acid substitutions (S/T → A, and S/T → D) that resemble the phosphorylation status similar to the phosphorylation status of MED1-IDR during Late-S phase. (D) Representative images of MCF10A cells expressing mNeonGreen fused to the wild-type MED1-IDR (NG-IDR^WT^) or mutated MED1-IDR variants (NG-IDR^G1^, NG-IDR^G1/S^, NG-IDR^Late-S^). (E) Line plot showing the relationship between the fraction of the cell population with mNeonGreen foci (>0.5 μm diameter) formation for the distinct mNeonGreen-MED1^IDR^ variants and the integrated mNeonGreen intensity. The integrated mNeonGreen intensity is binned. Each line represents a different mNeonGreen-MED1^IDR^ variant (G1, G1/S, and Late S). Points and error bars are the mean and standard error, respectively, of 3 independent experiments. (** p < 0.01, **** p < 0.0001). (F) Line plot showing the average number of mNeonGreen foci per cell for the distinct mNeonGreen-MED1^IDR^ variants as a function of the integrated mNeonGreen intensity. The integrated mNeonGreen intensity is binned. Each line represents a different mNeonGreen-MED1^IDR^ variant (G1, G1/S, and Late S). Points and error bars are the mean and standard error, respectively, of 3 independent experiments. (* p < 0.05, ** p < 0.01, **** p < 0.0001). (G) Line plots of the Net Charge Per Residue (NCPR) values for the wild-type MED1-IDR (IDR^WT^) or mutated MED1-IDR variants (IDR^G1^, IDR^G1/S^, IDR^Late-S^). Colored circles indicate the charge status of the bin, light blue for low basic (> 1), dark blue for high basic (> 3), light red for low acidic (< −1), dark red for high acidic (< −3).

To test the role of the phosphorylation code on condensate dynamics, we generated phosphorylation-mimicking mutations and phosphorylation-deficient mutations in the different domains matching the pattern in G1, the G1/S, and late S phase cells. For example, mutation of all phosphorylated S/T residues in PD 2 and PD 4 to glutamic acid and all phosphorylated S/T residues in PD 1, 3, and 5 to alanine was used to mimic the late S-specific phosphorylation state of MED1^IDR^ (Figure 5C). We made doxycycline-inducible hybrid mNeonGreen-MED1^IDR^ phosphorylation mutants and expressed these in MCF10A cells. After 48 hours of induction, the G1 (PD 4)-specific phosphorylation mutants could only form condensates at expression levels 60% greater than that needed of the WT-IDR-NG, suggesting the G1-specific phosphorylation state is inhibitory to phase separation (Figures 5D-5F). In striking contrast, the G1/S (PD 2)-specific phosphorylation mutations favored phase separation, doing so at a threshold half that of the G1 mutants and forming more condensates (Figure 5F). Importantly, this is consistent with the timing of MED1 condensate formation at the G1/S transition (Figure 1E). The late S phase-specific mutations behaved similar to the G1-mutants (Figures 5D-5F). Interestingly, the G1/S phosphorylation state created more prominent blocks of alternating charges (Figures 5G and S5C), in line with previous studies implicating charge blockiness as a determinant of phase separation[7, 8]. These results suggest the different phosphorylation clusters in MED1^IDR^ are major determinants of MED1 phase separation and collectively act as a code to specify the temporal dynamics of condensate formation and dissolution across S phase. Moreover, the phosphorylation code explains why ATR, CHK1, and CDK1/2 inhibitors alter MED1 condensates across S phase, as they are predicted to induce broad changes to MED1^IDR^ phosphorylation.

### Deregulation of MED1 causes ATR inhibitor-induced lethality

ATR is essential for the viability of proliferating cells[34–36], and accordingly, we asked whether the essentiality of ATR is linked to its role in regulating MED1 condensate dynamics. To this end, we first considered a published CRISPR-Cas9 screen looking for genetic drivers of lethality in ATR-inhibited cells[37]. We averaged the normalized Z-scores from this screen in three cell lines (MCF10A, HCT116, and HEK293) to identify genes whose protein products induce lethality in multiple cell lines with ATR inhibition. MED1 knockout was among the top hits that caused resistance to ATR inhibition (Figures 6A and S6A). Notably, examination of each mediator subunit indicated that loss of the mediator complex does not consistently influence the response to ATR inhibition, as the different subunits were distributed equally across the normalized Z-scores (Figure S6B). This suggests that ATR inhibition leads to MED1-induced lethality independently of a functional mediator complex.

**Figure 6.**
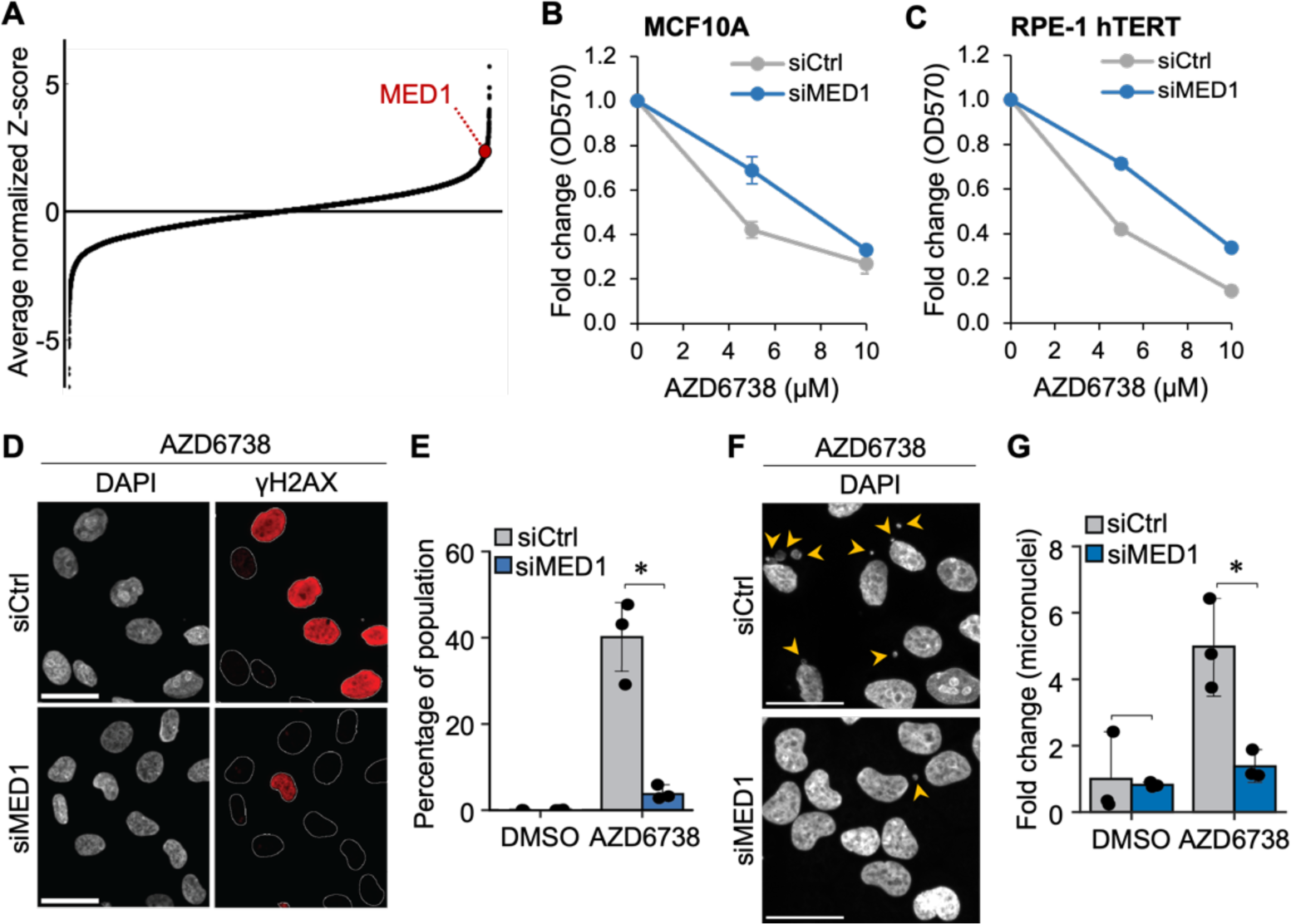
Deregulation of MED1 causes genome instability and loss of viability in ATR-inhibited cells. (A) Average normalized Z-score from a genome-wide CRISPR-Cas9 screen in ATR-inhibited cells[37]. MED1 is highlighted to show MED1 knockout increases survival of ATR-inhibited cells. Averages were calculated from the normalized Z-scores for MCF10A, HCT116, and HEK293 cells. (B) Cell viability assay of MCF10A cells transfected with non-targeting siRNA (siCtrl) and siRNA targeting MED1 (siMED1) and treated with 0, 5, or 10 μM AZD6738 for 48 h. (C) Cell viability assay of RPE-1 hTERT cells transfected with non-targeting siRNA (siCtrl) and siRNA targeting MED1 (siMED1) and treated with 0, 5, or 10 μM AZD6738 for 48 h. (D) Representative images of γH2AX and DAPI in MCF10A cells transfected with siCtrl or siMED1 and treated with 5 μM AZD6738 for 16 h. Scale bar, 30 μm. (E) Percent of late S and G2 phase cells with pan-nuclear γH2AX staining. Cells were treated as in (D). Bars represent the mean of 3 independent experiments (each experiment shown as a single dot) and whiskers indicate the standard error of the mean. (* p < 0.05) (F) Representative images of DAPI in cells treated as in (D). Yellow arrows mark the micronuclei. Scale bar, 30 μm. (G) Fold-change in micronuclei/cell treated as in (D). Bars represent the mean of 3 independent experiments (each experiment shown as a single dot) and whiskers indicate the standard error of the mean. (* p < 0.05)

To confirm that MED1 induces cell death in ATR-inhibited cells, we silenced MED1 expression with pooled siRNAs (siMED1) in MCF10A and RPE-1 hTERT cells and inhibited ATR with AZD6738. After 48 hours of inhibition, cells were plated as single cells and allowed to form colonies in normal, untreated growth medium for 10 days. In the non-targeting siCTRL-transfected cells, ATR inhibition resulted in a significant reduction in colony formation. Importantly, MED1 knockdown rescued survival approximately 2-fold (Figures 6B and 6C), confirming MED1 as a driver of lethality in ATR-inhibited cells.

ATR inhibitors, either used alone or in combination with a replication stressor, induce pan-nuclear γH2AX staining, indicative of genome-wide DNA damage[21, 38]. To determine if the DNA damage is MED1-dependent, we again silenced MED1 with pooled siRNAs and inhibited ATR with AZD6738. Approximately 40% of siCTRL cells in late S and G2 cell cycle phases exhibited pan-nuclear DNA damage after 16 h of ATR inhibition (Figures 6D and E). Strikingly, MED1 knockdown caused a near complete rescue of ATR inhibition-induced DNA damage. In addition, MED1 knockdown significantly reduced micronucleus formation in ATR-inhibited cells (Figures 6F and G). These data suggest ATR inhibition causes deregulation of MED1 leading to global DNA damage, chromosome instability, and loss of cell viability.

### ATR ensures linker histone H1 duplication and genome stability

Given that ATR controls MED1 condensate dynamics at HLBs in S phase and ATR inhibition results in MED1-dependent DNA damage, we reasoned the two phenomena may be linked. HLBs form around the 3 replication-dependent histone gene clusters, the largest of which, HIST1 located on human chromosome 6p21-p22, encodes 12 H2A genes, 15 H2B genes, 10 H3 genes, 12 H4 genes, and 1 gene each of the linker H1 genes (H1.1, H1.2, H1.3, H1.4, H1.5, H1.t)[39]. HLBs couple histone expression to the S phase to ensure that cells duplicate their histone content concomitantly with DNA duplication[25]. If the MED1-induced pan-nuclear DNA damage is due to the deregulation of MED1 condensates at HLBs, then ATR inhibition should result in a detrimental change in histone expression. To test this, we imaged the replication-dependent histones, and using QIBC, confirmed that the histones were, for the most part, duplicated across S phase (Figures S7A and S7B). We inhibited ATR for 16 hours to allow a significant percent of the population to pass through S phase and then imaged each of the core histones and linker histones. We observed a moderate, but consistent increase in most histones across the cell cycle when ATR was inhibited (Figure S7B). We co-stained cells with γH2AX to determine if changes in histone duplication correlated with ATRi-induced pan-nuclear DNA damage. Strikingly, cells that exhibited pan-nuclear DNA damage failed to duplicate their linker histone H1.1 (Figures 7A-7D). Cells also failed to duplicate linker histone H1.3, though the effects on H1.3 were mild (Figure S7B). Importantly, CHK1 inhibition also caused a failure to duplicate H1.1 in cells with pan-nuclear DNA damage (Figure 7E), further strengthening the link between deregulated MED1 condensates and DNA damage.

**Figure 7.**
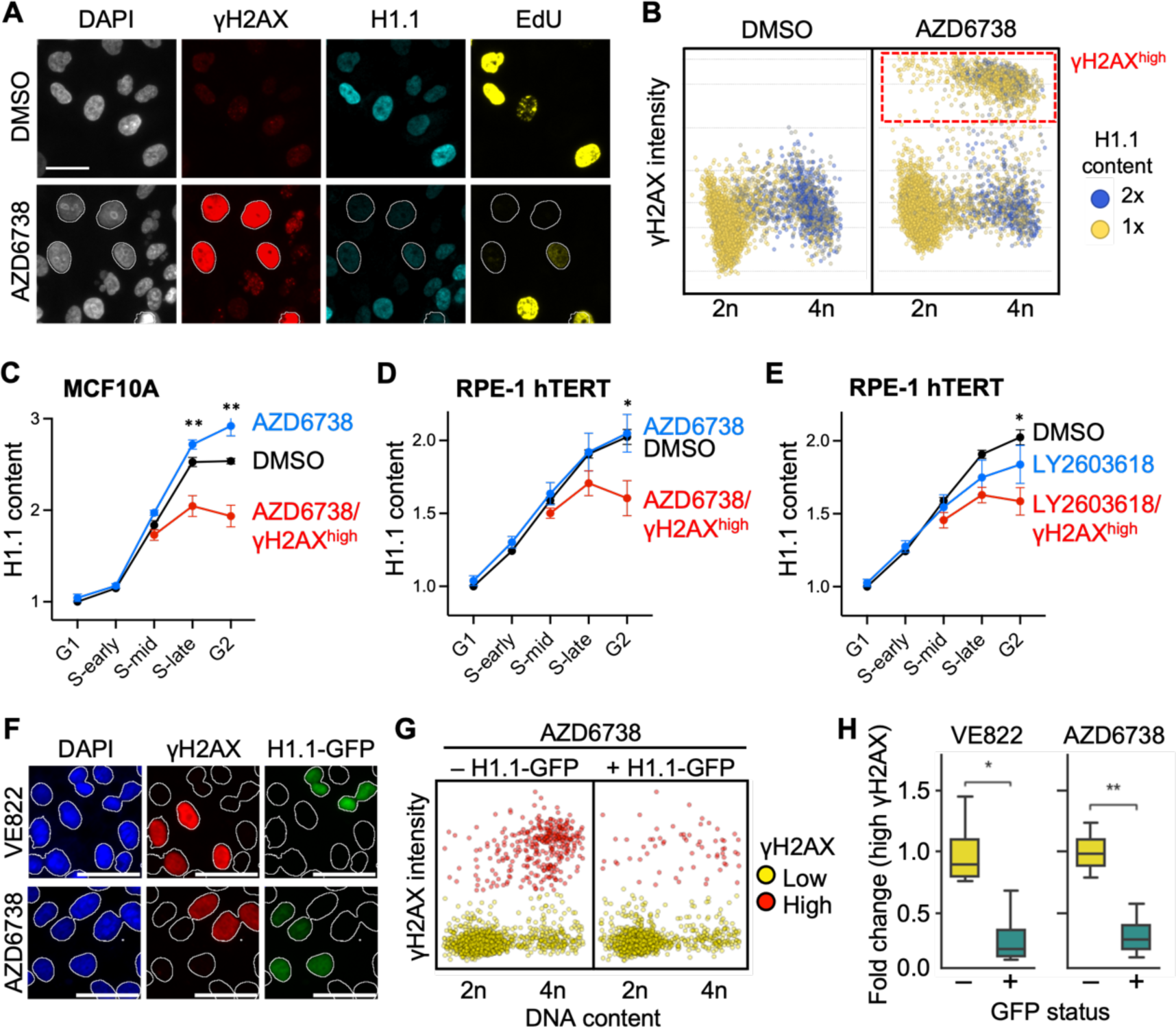
Defective linker histone H1.1 duplication causes pan-nuclear DNA damage in ATR-inhibited cells. (A) Representative images of DAPI, γH2AX, H1.1, and EdU in MCF10A cells treated with DMSO or 5 μM AZD6738 for 16 h. Nuclear outlines mark the cells with pan-nuclear γH2AX. Scale bar, 30 μm. (B) Scatter plots of cells treated as in (A) showing DAPI integrated intensity (linear scale) vs. γH2AX mean intensity (log_2_ scale). Red box indicates cells with pan-nuclear γH2AX. Color scale indicates H1.1 integrated intensity with H1.1 levels doubling across S phase. (C) Line plots of H1.1 content (integrated intensity relative to G1 phase population average) in each cell cycle phase of MCF10A cells. Cells were treated as in (A). Red line shows H1.1 levels in the cells with pan-nuclear γH2AX (γH2AX^high^). (** p < 0.01) (D) Line plots of H1.1 content as described in (C) but in RPE-1 hTERT cells. (* p < 0.05) (E) Line plots of H1.1 content as described in (C) but in RPE-1 hTERT cells treated with DMSO or 2 μM LY2603618 (CHK1 inhibitor) for 16 h. (* p < 0.05) (F) Representative images of DAPI, γH2AX, and H1.1-GFP in MCF10A cells treated with 5 μM AZD6738 or 5 μM VE822 for 16 h. Scale bar, 30 μm. (G) Scatter plots of cells treated as in (F). Plots show γH2AX in cells either negative for H1.1-GFP (left) or positive for H1.1-GFP (right) and are colored to indicate pan-nuclear γH2AX (red) or normal γH2AX (yellow). (H) Boxplots showing the fold change of pan-nuclear γH2AX positive cells of MCF10A cells positive (+) or negative (–) for H1.1-GFP expression. Plots are divided by ATR inhibition treatment, 5 μM VE822 and 5 μM AZD6738. (* p < 0.05, ** p < 0.005)

To determine if the shortage of linker histones caused the global DNA damage, we rescued H1.1 levels by ectopic expression of H1.1-GFP and measured γH2AX levels following ATR inhibition (Figures 7F and S7C). The percentage of cells with pan-nuclear DNA damage upon ATR inhibition in late S and G2 phases was reduced by 50% in H1.1-GFP positive cells compared to H1.1-GFP negative cells (Figures 7G and 7H). These data are strong evidence that pan-nuclear DNA damage in ATR-inhibited cells arises as a consequence of failed duplication of linker histone H1.1 and perhaps H1.3 as cells progress through S phase. Combined with the MED1 knockdown data, our results point to ATR-dependent regulation of transcription condensates in S phase via a phosphorylation code as a key regulator of histone duplication and genome integrity.

## DISCUSSION

Our study uncovers the dynamic nature of large transcription condensates that appear precisely at the G1/S cell cycle transition and dissolve during S phase progression. Their dynamics are largely dictated by a phosphorylation code present in the MED^IDR^. While phosphorylation has been shown to regulate condensate formation in certain contexts and dissolution in others[8, 9, 40], to our knowledge this is the first example of a phosphorylation code acting on an IDR.

An important distinction of the MED1^IDR^ phosphorylation code from other phosphorylation events regulating condensates is that it is the collective contribution of multisite phosphorylation within a single cluster or combination of clusters that regulates MED^IDR^ dynamics. We speculate the different phosphorylation clusters that make up the code alter the pattern and distribution of charged amino acids in the MED^IDR^. In this way, one cluster may enhance the alternating charged blocks favoring the formation of a compartment, whereas another cluster may neutralize a charged block to dissolve the compartment. Given that many IDRs are enriched with S/T residues[33], it is tempting to speculate that phosphorylation codes made up of clusters of phosphorylation events may be a general regulatory mechanism for condensate dynamics.

Several candidate kinases could potentially phosphorylate MED1^IDR^, including ATR, CHK1, and CDK1/2. These kinases are highly active in S phase, and importantly, many of the clustered phosphorylation events occur within the motifs they recognize. ATR and CHK1 are known to suppress CDK1/2 activity[13, 17, 27, 29], and thus, inhibiting ATR and CHK1 would likely cause broad changes to the phosphorylation pattern on MED^IDR^. Indeed, ATR and CHK1 inhibition causes a delayed dissolution of condensates by altering the electrostatic properties of MED1. This is consistent with the hypothesis that phosphorylation clusters either create or neutralize alternating charged blocks in MED1^IDR^.

Large, diffraction-sized condensates of RNAPII and Mediator have previously been observed in mouse embryonic stem cells and are thought to bring super-enhancers in close proximity to promoters to drive gene expression[3]. Recently, it was shown that a large condensate forms near the super-enhancer controlled Sox2 gene in mouse embryonic stem cells. Transient 3-way interactions involving the condensate, the Sox2 gene, and its super-enhancer correlated with a burst in transcription of Sox2, suggesting the condensate regulates gene expression[41]. How prevalent these large condensates are and what transcriptional programs they control remains unknown. Moreover, it is unknown if these large transcription condensates are unique features of embryonic stem cells or if they are also common and functional in other cell types.

The large transcription condensates that appear at the G1/S transition form at HLBs. The timing of their appearance ensures the expression of replication-dependent histone genes is up-regulated in S phase. HLBs are compartments that exist stably throughout interphase. The transient formation of MED1 condensates within these nuclear bodies suggests that transcription condensates can form and dissolve, not just as their own entity, but as part of larger, stable compartments. Notably, the dissolution of the condensate in mid-S phase suggests transcription of histone genes occurs primarily in early S phase. This, perhaps, prevents over-expression of histones in late S phase and/or unwanted histone biosynthesis in the G2 phase. Indeed, evidence suggests cells tightly control histone level, with overexpression linked to cell toxicity[42].

Inhibition of either ATR or CHK1 resulted in an increase in most histones across the cell cycle, with the exception being the linker histones H1.1 and H1.3. H1.1 and H1.3 were reduced in late S phase and the G2 phase causing extensive DNA damage throughout the nucleus. It is unknown why these two histones were uniquely affected by ATR inhibition, but it may be that compensatory mechanisms decrease histone levels when they exceed a threshold and this compensation more adversely affects H1.1 and H1.3. It is also unknown why a decrease in H1.1 and H1.3 would lead to widespread DNA damage. In general, linker histones promote chromatin compaction[43] and this, in turn, affects transcription and replication. We speculate H1.1 and H1.3 may act to keep specific regions in a more densely-packed state, and that loss of these linker histones cause an aberrant opening of chromatin leading to replication-associated DNA damage. Alternatively, H1.1 and H1.3 may have unique functions independent of chromatin compaction. Indeed, the different linker histones appear to have distinct functions including regulation of origin firing and DNA break repair[44, 45]. To date, little is known about the distinct functions of H1.1 and H1.3, and thus, further research is needed to determine why their loss leads to global DNA damage. Nonetheless, the decrease in H1.1 and H1.3 creates an imbalance in the pool of linker histones, a phenotype that has been linked to genome instability, cGAS activation, and immune dysfunction in human disease[46].

An important finding of this study involves the enrichment of ATR within HLBs. ATR accumulates in this compartment during S phase and signals the dissolution of MED1. While MED1 exhibits liquid droplet-like properties as suggested by its disruption by 1,6-HXD, ATR does not. In clear contrast to MED1, ATR foci are enlarged upon 1,6-HXD treatment, leaving the nature of ATR enrichment in HLBs unknown. As part of the DNA damage response (DDR), ATR is recruited to DNA damage foci through binding to RPA-coated single-stranded DNA[47]. Moreover, optogenetic studies have linked the DDR to condensate formation and ATR activation[40, 48, 49]. However, we have not been able to detect RPA at HLBs and ATR signaling at HLBs does not activate the DDR, suggesting ATR recruitment to these compartments involves a novel, unknown, DDR-independent mechanism. Despite the absence of RPA, ATR is active within HLBs and signals downstream suppression of CDK1 to prevent FOXM1 phosphorylation in S phase as part of the recently described S/G2 checkpoint[13, 27, 29]. Given that ATR coordinates histone synthesis with S phase and suppresses G2/M transcription in S phase within HLBs, we propose this compartment to be a critical hub of S/G2 checkpoint signaling.

The S phase is a period of rapid change to chromatin and without the precise coordination of replication and transcription, genome-wide DNA damage would lead to loss of viability. This coordination is achieved, in part, through the compartmentalization of replication and transcription into spatially-distinct regions. These compartments must be dynamic to account for their spatiotemporal patterns across S phase, and importantly, the ATR pathway and the MED1^IDR^ phosphorylation code provides an elegant means to do so.

## ACKNOWLEDGEMENTS

We thank members of the Saldivar laboratory for helpful discussions on the work presented in this paper. C.O.M. and J.L. are funded by an award from the Cancer Early Detection Advanced Research Center (CEDAR). C.S. is funded by a National Institutes of Health (NIH) Institutional Training Grant T32GM142619. Work in the Hamperl laboratory is funded by the Helmholtz Association, the German Research Foundation (DFG) Project-ID 213249687 (SFB 1064) and the European Research Council (ERC starting grant 852798). J.C.S. is funded by CEDAR, an NIH grant GM147710, and by a Career Development Award to Promote Diversity and Inclusion from the American Association for Cancer Research (AACR) 21-20-26-SALD.

## AUTHOR CONTRIBUTIONS

C.O.M. and J.C.S. conceived the project, conducted experiments, analyzed data, prepared figures, and wrote the manuscript. C.S., P.R., M.W., and S.H. performed several key experiments. J.L. helped analyze the data.

## METHODS

### Cell lines and reagents

MCF10A and RPE-1 hTERT cells were purchased from ATCC. MCF10A cells were cultured in DMEM:F12 (Thermo Fisher Scientific, 11320082) base medium supplemented with 5% horse serum (Thermo Fisher Scientific, 16050122), 20ng/ml EGF (Peprotech, AF-100-15), 10 ug/ml insulin (Millipore Sigma, I-1882), 0.5 mg/ml hydrocortisone (Millipore Sigma, H-0888), 100 ng/ml cholera toxin (Millipore Sigma, C-8052), and 1x penicillin/streptomycin (Thermo Fisher Scientific, 15140122). RPE-1 hTERT were cultured in DMEM:F12 base medium supplemented with 10% fetal bovine serum (Thermo Fisher Scientific, A5256801), 0.1 mg/ml hygromycin B (Millipore Sigma, 10843555001), and 1x penicillin/streptomycin.

AZD6738 (Selleck Chemicals, S7693) and VE-822 (Selleck Chemicals, S7102) were used as ATR inhibitors from a stock concentration of 10 mg/ml. Additional inhibitors used in the study were LY2603618 (Selleck Chemicals, S2626), CHIR-124 (Selleck Chemicals, S2683), RO3306 (Selleck Chemicals, S7747), BMS-265246 (Selleck Chemicals, S2014), and NU6140 (Thermo Fisher Scientific, 33-011-0). 1,6 hexanediol (Sigma-Aldrich, 240117-50G) was diluted to a 50% w/v concentration in water.

### Antibodies

The antibodies used in Immunofluorescence include: anti-phospho-Histone H2A.X (1:1000, Millipore Sigma, 05-636-I), anti-phospho-ATR (1:500, Cell Signaling Technology, 2853S), anti-TRAP220/MED1 (1:1000, Abcam, ab64965), anti-RNAPOLII (1:1000, Abcam, ab26721), anti-NPAT (1:1000, BD Biosciences, 611344), anti-phospho-FOXM1 (1:250, Cell Signaling Technology, 14655S), anti-BRD4 (1:500, Abcam, ab128874), anti-RPA32/RPA2 (1:500, Abcam, ab76420), anti-Coilin (1:1000, Cell Signaling Technology, 14168T), anti-53BP1 (Abcam, ab237174), anti-H1.1 (1:1000, Abcam, ab254394), anti-H1.2 (1:500, GeneTex, GTX122561), anti-H1.3 (1:1000, Abcam, ab183736), anti-H1.4 (1:500, Cell Signaling Technology, 41328), anti-H1.5 (1:1000, Abcam, ab18208), anti-H2A (1:500, Cell Signaling Technology, 12349), anti-H2B (1:500, Cell Signaling Technology, 12364), anti-H3 (1:500, Cell Signaling Technology, 4499S), anti-H4(1:500, Cell Signaling Technology, 13919).

### Plasmid transfection

MCF10A cells were seeded at 8,000 cells per well in glass-bottom 96-well plates (Cellvis P96-1.5P). 24 h post-seeding, cells were transfected with pEGFP-H1.1 plasmid (Addgene, 32894) using Fugene 6 transfection reagent (Promega, E2691), following manufacturer’s guidelines. The medium was replaced with fresh medium 24 h post transfection. Cells were analyzed 48 h post-transfection.

### siRNA transfection

Cells were seeded at 8,000 cells per well in glass-bottom 96-well plates (Cellvis P96-1.5P) and reverse-transfected with siRNAs at 20 nM using DharmaFECT 1 (Horizon Discovery T-2005-01) following manufacturer’s guidelines. 16 h post-transfection, media was changed and replaced with normal growth media. Cells were incubated for a total of 40 h or 48 h post-transfection. In experiments involving a 16-hour treatment with ATR inhibitors, cells were reverse-transfected with siRNA, then after 32 h incubation, media was replaced with media containing either DMSO (mock treatment) or ATR inhibitors. Cells were analyzed 48 h post-transfection.

The siRNA used include MED1 siRNA (Horizon Discovery, L-004126-00-0005), ATR siRNA (Horizon Discovery, J-003202-20-0002), and control (Non-targeting Pool) siRNA (Horizon Discovery, D-001810-10-05).

### Immunostaining

Cell samples were fixed with 4% paraformaldehyde (PFA) diluted in phosphate-buffered saline (PBS) for 10 min, permeabilized with ice-cold methanol for 10 min, washed 3 times in 1X PBS, and blocked in 1% bovine serum albumin (BSA) in PBS for 30 min at room temperature. For 5-ethynyl-2’-deoxyuridine (EdU) staining, the Click-iT reaction was carried out following permeabilization using the Click-iT Cell Reaction Buffer kit (Thermo Fisher Scientific C10269) and Alexa Fluor™ 647 Azide (Thermo Fisher Scientific A10277) according to the manufacturer’s guidelines. Specifically, cells were washed with 3% BSA/PBS and incubated with the Click-iT reaction mixture with 1 µg/ml of Alexa Fluor™ 647 Azide for 30 min with at room temperature on a bench rocker. After Click-iT reaction mixture incubation, cells were washed with 3% BSA/PBS, washed 3 times in 1X PBS, and blocked in 1% BSA in PBS for 30 min at room temperature. Following blocking of samples, the cells were incubated with primary antibodies. The primary antibodies were diluted in 1% BSA/PBS and incubated overnight with constant agitation at 4°C. Cells were then washed 3 times in 1X PBS and co-stained with DAPI (5 μg/mL) and secondary antibodies (1:1000), anti-rabbit Alexa Fluor 488 conjugated antibody (Thermo Fisher Scientific A-11008) and anti-mouse Alexa Fluor 568 conjugated (Thermo Fisher Scientific A-11004), diluted in 1% BSA and incubated for 1 h at room temperature on a bench rocker. Cells were washed 3 times in 1X PBS before imaging.

### Quantitative image-based cytometry and analysis

Cells were grown in glass-bottom 96-well plates (Cellvis P96-1.5P) and imaged on a fully automated ImageXpress Micro (Molecular Devices) at 20X and 40X. Image analysis was performed using CellProfiler^1^ (Broad Institute). Intensity measurements were within a nuclear mask generated from DAPI-stained images. Background levels were determined for each fluorescence marker by assessing histogram plots of the signal intensities in each pixel across several images. The average background value across these images was then subtracted from the mean intensity of the given fluorescent marker in each cell. DNA content was determined using DAPI integrated intensity. Identification and quantification of foci (ATR foci, MED1 foci, and NPAT foci) was performed using an *in-house* developed CellProfiler pipeline. Identification and quantification of micronuclei were derived from the analysis of DAPI-stained images.

### Cell viability assay

MCF10A and RPE1 cells were seeded at 50,000 cells per well in 12-well plates (Corning Costar 3513) and reverse-transfected with siRNA (siMED1 or siControl) at 50 nM using DharmaFECT 1 (Horizon Discovery T-2005-01) as per the manufacturer’s guidelines. 16 h post-transfection, media was changed and replaced with media containing either DMSO (mock treatment) or AZD6738 (5 μM). After an initial 24-hour treatment period, the media was refreshed once again with the respective treatment media (DMSO or AZD6738). Following a cumulative 48 h treatment, cells were harvested using TrypLE (Thermo Fisher Scientific 12605010) and re-seeded at 500 cells per well in 6-well plates using untreated media. Cells were subsequently cultured over a 7-day period with media changes occurring every 2 days. Upon reaching the 7-day mark, cells were rinsed once with PBS, and once with ddH2O, before being fixed with 1 ml per well of 0.5% crystal violet staining solution (0.5 g crystal violet powder [Milipore Sigma C0775], 80 ml ddH2O, and 20 ml methanol) incubate for 20 min at room temperature on a bench rocker. Post-incubation, samples were washed 4X with ddH2O. The plates were then air-dried for 18 h, after which, cells were incubated in methanol for 20 min at room temperature on a bench rocker. Samples were transfer to a 96 well plate using 100 ul per well. The optical density at 570 nm (OD_570_) was measured in triplicate for each well utilizing a Tecan’s Spark® 20M plate reader. Data represent the mean and the standard error of the mean for three independent experiments.

### Proximity ligation assay

All steps were conducted at room temperature unless stated otherwise. Cells were seeded in a 96-well imaging plate (Ibidi, 89626) and treated with 10 μM EdU for 30 min before pre-extraction. Pre-extraction was done with 0.5 % Triton X-100 in PBS for 2 min. Next, cells were washed with PBS and fixed with 4 % PFA in PBS for 15 min. Following fixation, cells were washed with PBS wash, permeabilized with 0.2 % Triton-X in PBS for 4 min and washed with PBS again. EdU click-it reaction solution (100 mM Tris-HCl pH 8.5, 1 mM CuSO_4_, 100 mM Ascorbic acid, 0.9 µg Alexa Fluor 594) was added, and cells were incubated in the dark for 30 min. Cells were then blocked in 5 % BSA in PBS for 45 min, followed by application of primary antibodies against ATR (1:300, Cell Signaling Technolgy, 13934) and RNAPII (1:2000, Merck Millipore, 05-623) diluted in 5 % BSA in PBS. Cells were incubated at 4 °C overnight. The next day, cells were washed twice with PBS. Subsequently, Duolink PLUS (Sigma Aldrich, DUO92001) and MINUS (Sigma Aldrich, DUO92005) probes diluted with Duolink Antibody Diluent (1:10) were applied for 1 h at 37 °C. The cells were washed twice with Wash buffer A (Sigma Aldrich, DUO82049) and the ligation solution (1x Duolink ligation buffer, Ligase at 1:70 dilution) (Sigma Aldrich, DUO92014) was applied for 30 min at 37 °C. After two additional washes with Wash buffer A the amplification solution (1x Amplification buffer, Polymerase at 1:140 dilution) (Sigma Aldrich, DUO92014) was added and cells were incubated in the dark at 37°C for 100 min. The cells were then washed twice with Wash buffer B (Sigma Aldrich, DUO82049) and stained with DAPI in 3 % BSA in PBS for 1 h. Finally, cells were washed twice with PBS and stored at 4 °C until imaging. Images were acquired on a Nikon T2 inverted microscope equipped with an Andor Dragonfly spinning disk, a 40X air objective and an iXon Life 888 EMCCD camera. Per condition 81 positions were imaged at which a Z-stack of 7 images across 10 µm was acquired. Image analysis was conducted as described previously[50].

### mNeonGreen-MED1^IDR^ construction

The vectors containing mNeonGreen-MED1^IDR^ (NG-IDR) variants were constructed using a lentiviral vector backbone, LTO-NG vector, with the mNeonGreen gene under the control of a doxycycline inducible promoter (Tet-ON), and the rtTA3 gene fused to the blasticidin resistant marker (BlastR) by a 2A self-cleavage peptide (P2A) under the control of a SV40 promoter. The doxycycline inducible promoter was obtained from the pCW57.1 plasmid (Addgene, 41393), the mNeonGreen gene was obtained from the mNeonGreen-mTurquoise2 plasmid (Addgene, 98886), the rtTA3 gene fused to the blasticidin resistant marker (BlastR) was adapted from the pLenti CMV rtTA3 Blast (w756-1) plasmid (Addgene, 26429).

The vector containing MED1^IDR^ wild-type (NG-IDR^WT^) was constructed by PCR amplification of the sequence comprising the amino acids 560 to 1582 of MED1, utilizing the plasmid pWZL hygro Flag HA TRAP220 wt (Addgene, 17433). The amplified MED1^IDR^ sequence was inserted into the LTO-NG vector backbone using Golden Gate Assembly, resulting in a plasmid used for the expression of NG-IDR^WT^.

Subsequently, the MED1^IDR^ was divided into 5 sequence blocks, each containing a phospho-domain. For each sequence block, we generated an OFF version, in which the sequences for all the serine / threonine residues identified within the phospho-domain were mutated to generated a substitution for alanine, and a ON version, in which the serine / threonine positions were mutated to aspartic acid. The version of the 5 sequence blocks was synthesized using g-Blocks (Integrated DNA Technologies). The g-blocks also contained compatible overhangs which allow the assembly of any combination of sequence block with the LTO-NG vector backbone via Golden Gate Assembly.

### Lentivirus production and transductions

Second generation lentiviruses were generated using HEK293T cells, purchased from ATCC. Cells were transfected using FuGENE 6 transfection reagent (Promega, E2691). Briefly, a transfection mixture was prepared by diluting 6 μg of the mNeonGreen-MED1^IDR^ variant vector, 6 μg of the psPAX2 plasmid (Addgene plasmid no. 12260) and 0.6 μg of the pMD2.g plasmid (Addgene, 12259) in Opti-MEM medium (Thermo Fisher Scientific, 31985062) and mixing with 36 μl FuGENE 6 transfection reagent according to the manufacturer’s instructions. The transfection mixture was then added directly into a 100 mm dish, followed by seeding 1×10^6^ cells and a gentle swirling to ensure uniform distribution. The medium was replaced with fresh medium 8 h post transfection and the virus-containing medium was collected after 48 h. The virus was concentrated using a Lenti-X concentrator (Clontech, 631232) according to the manufacturer’s protocol.

Cell transduction was performed by seeding MCF10A cells onto 12-well plates at a density of 1×10^5^ cells per well. After 24 h, the medium was replaced with medium containing 2X concentrated virus medium and 8 μg/ml polybrene (Millipore Sigma, TR-1003-G). The virus-containing medium was replaced with fresh medium 24 h post transduction.

### Net Charge Per Residue (NCPR) calculation

The Net Charge Per Residue (NCPR) calculation was performed following the protocol outlined previously[7]. Briefly, NCPR was determined using a sliding window approach encompassing 10 amino acids. Specifically, a sliding window was identified as a low acidic or basic block if its NCPR value exceeded the absolute threshold of 1, high acidic or basic block if its NCPR value exceeded the absolute threshold of 3, and with negativity indicating acidity and positivity indicating basicity.

### Statistical analyses

Statistical metrics encompassing the number of biological replicates (n), standard deviation and statistical significance are reported in both the visuals graphs and the accompanying figure captions. The threshold for determining statistical significance has been established at a p-value less than 0.05, assessed using two-tailed Student’s t-test or One-Way ANOVA test, contingent on the specific requirements of the data set under evaluation.

## SUPPLEMENTAL INFORMATION

### Supplemental Figures

**Figure S1.**
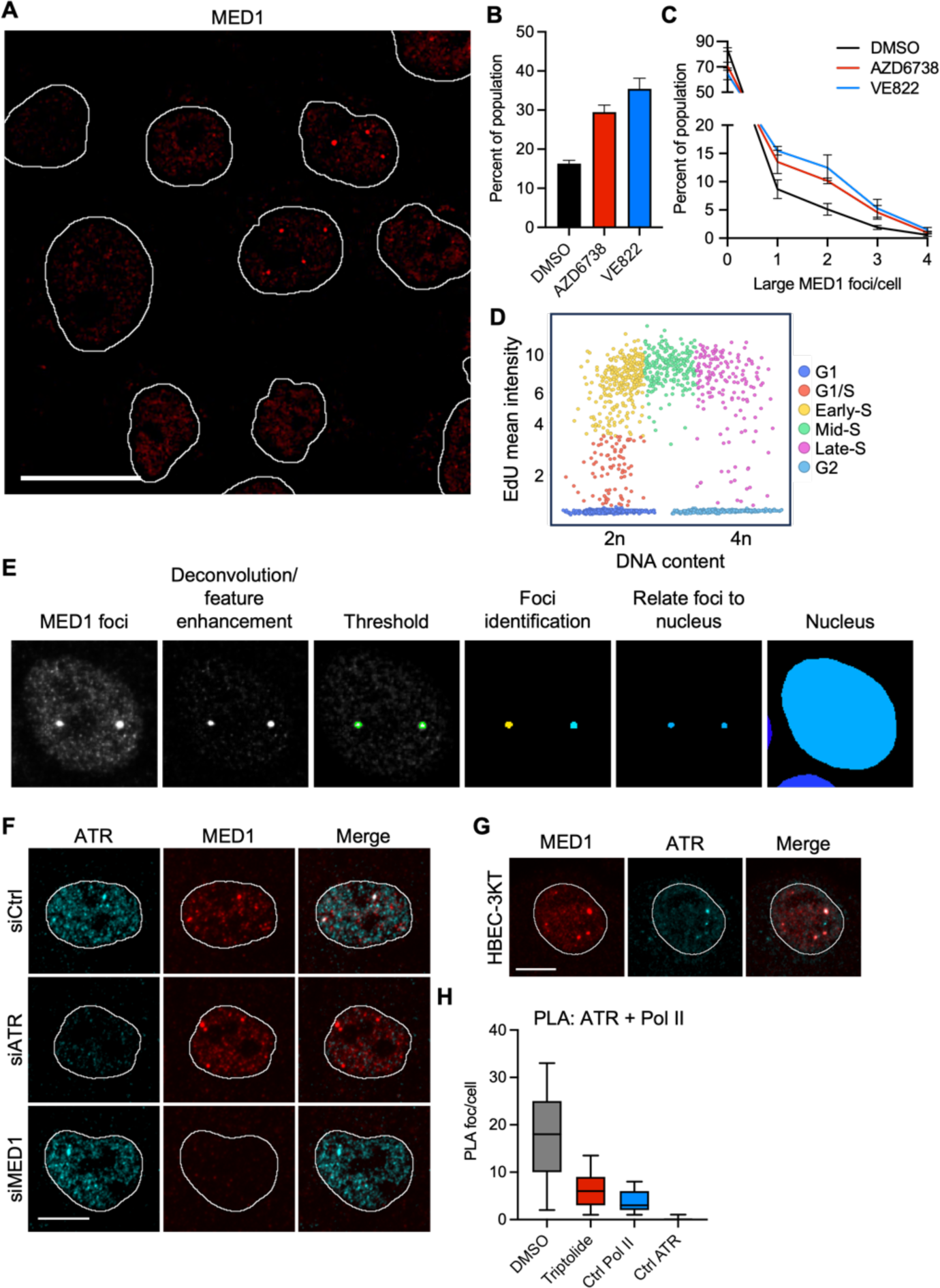
ATR colocalizes with MED1 in multiple cell lines and cell types. (A) Representative image of MED1 in MCF10A cells showing large condensates in only a subset of cells. Scale bar 20 μm. (B) Quantification of (A) showing the percent of cells with large MED1 foci. Mean and standard error 3 independent experiments. (C) Further quantification of (B) showing the percent of cells with 0-4 large MED1 foci. Mean and standard error of 3 independent experiments. (D) Scatterplot of DNA content (DAPI integrated intensity) vs. EdU mean intensity. Cells are colored by their respective cell cycle phase or sub-phase. (E) Representative image showing a cell with large MED1 foci and image processing steps to identify foci and match to parent nucleus. (F) Representative images of ATR and MED1 in MCF10A cells transfected with non-targeting control, MED1, or ATR siRNAs. Scale bar 10 μm. (G) Representative images of MED and ATR in immortalized human bronchiolar epithelial cells (HBEC-3KT). Scale bar 10 μm. (H) Boxplots showing the quantification of PLA foci of ATR and RNAPII per cell. Cells were treated with mock (DMSO) or 1 μM Triptolide for 2 h. Ctrl Pol II and Ctrl ATR are single-antibody controls from cells treated with DMSO for 1h.

**Figure S2.**
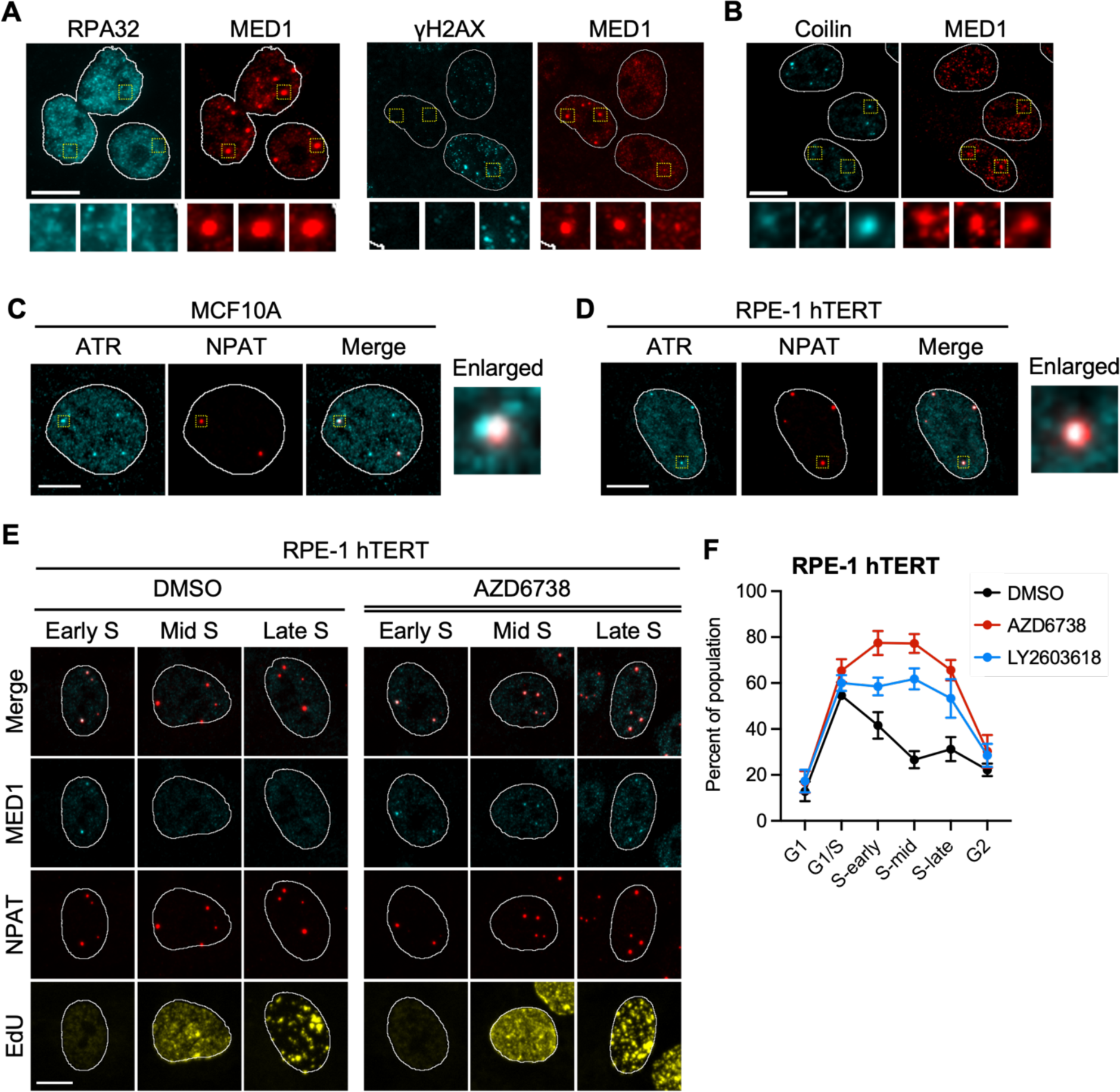
Large ATR-MED1 condensates mark the histone locus body. (A) Representative images showing MED1 and yH2AX, MED1 and RPA32, and MED1 and 53BP1 foci in MCF10A cells. Scale bar 10 μm. (B) Representative images showing MED1 and coilin foci in MCF10A cells. Scale bar 10 μm. (C) Representative images showing ATR and NPAT foci colocalization in MCF10A cells. Scale bar 10 μm. (D) Representative images showing ATR and NPAT foci colocalization in RPE-1 hTERT cells. Scale bar 10 μm. (E) Representative images of MED1, NPAT and EdU in early, mid, and late S phase cells mock-treated (DMSO) or treated with 5 μM AZD6738 for 1 h. Scale bar 10 μm. (F) Line plots showing the percent of RPE-1 hTERT cells with large MED1 foci for each cell cycle phase. Cells were treated as in (E). Points and error bars are the mean and standard error, respectively, of 3 independent experiments.

**Figure S3.**
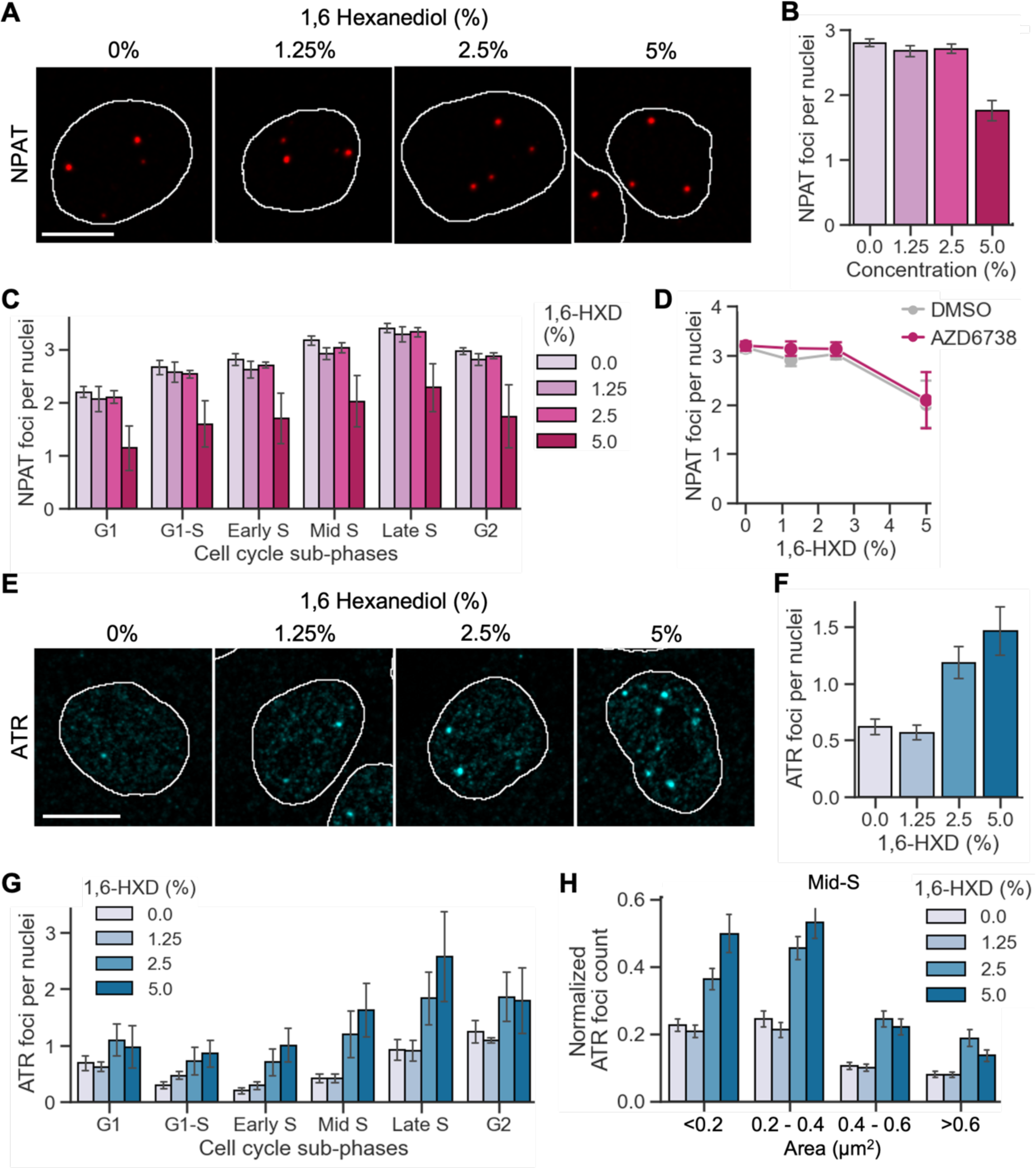
Effect of 1,6 hexanediol on NPAT and ATR foci. (A) Representative images of NPAT in mid-S phase MCF10A cells. Cells were treated with 0, 1.25, 2.5, or 5 % 1,6 hexanediol for 1 min. Scale bar, 10 μm. (B) Bar plot showing the quantification of NPAT foci in cells treated with varying concentrations (0%, 1.25%, 2.5%, and 5%) of 1,6-hexanediol (1,6-HXD) for 1 min. Bars and error bars are the mean and standard error, respectively, of 3 independent experiments. (C) Bar plot showing the quantification of NPAT foci in cells treated with varying concentrations (0%, 1.25%, 2.5%, and 5%) of 1,6-HXD for 1 min at different stages of the cell cycle. The color of each bar corresponds to the concentration of 1,6-HXD used. Bars and error bars are the mean and standard error, respectively, of 3 independent experiments. (D) Line plot showing the quantification of NPAT foci in cells treated with DMSO or 5 μM AZD6738 for 1 h, followed by a 1 min treatment with 0, 1.25, 2.5, and 5 % 1,6-HXD. Each line represents a different treatment condition (DMSO or AZD6738). Points and error bars are the mean and standard error, respectively, of 3 independent experiments. (E) Representative images of ATR in mid-S phase MCF10A cells. Cells were treated as in (A). Scale bar, 10 μm. (F) Bar plot showing the quantification of ATR foci in cells treated with varying concentrations (0%, 1.25%, 2.5%, and 5%) of 1,6-hexanediol (1,6-HXD) for 1 min. Bars and error bars are the mean and standard error, respectively, of 3 independent experiments. (G) Bar plot showing the quantification of ATR foci in cells treated with varying concentrations (0%, 1.25%, 2.5%, and 5%) of 1,6-HXD for 1 min at different stages of the cell cycle. The color of each bar corresponds to the concentration of 1,6-HXD used. Bars and error bars are the mean and standard error, respectively, of 3 independent experiments. (H) Bar plot illustrating the number of ATR foci per nuclei following a 1 min treatment with 0, 1.25, 2.5, and 5 % 1,6-HXD; across four distinct groups of ATR foci categorized by area: <0.2, 0.2 to 0.4, 0.4 to 0.6, and >0.6 µm². The color of each bar corresponds to the concentration of 1,6-HXD used. Bars and error bars are the mean and standard error, respectively, of 3 independent experiments.

**Figure S4.**
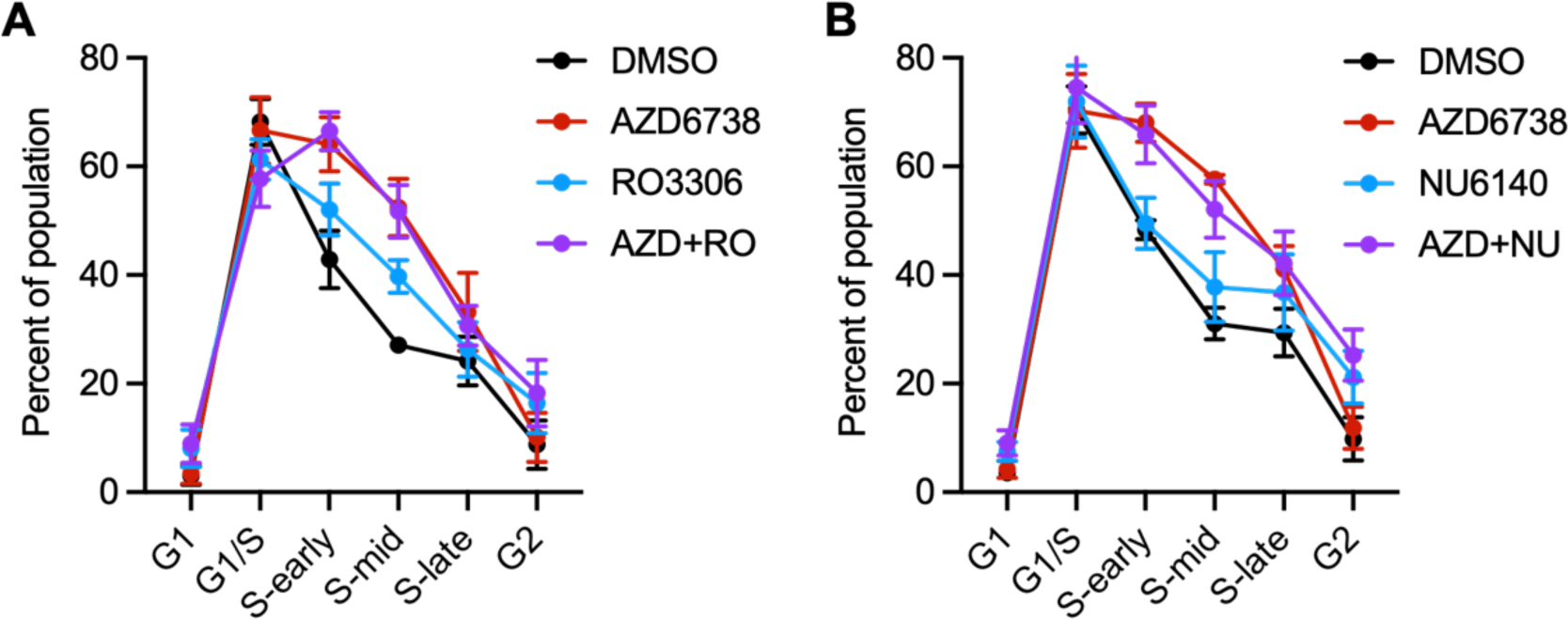
Effect of CDK2 inhibition or CDK1 inhibition on large MED1 condensates. (A) Line plots showing the percent of MCF10A cells with large MED1 foci for each cell cycle phase. Cells were mock-treated (DMSO) or treated with 5 μM AZD6738 and 5 μM RO-3306 individually or in combination for 1 h. Points and error bars are the mean and standard error, respectively, of 3 independent experiments. (B) Line plots showing the percent of MCF10A cells with large MED1 foci for each cell cycle phase. Cells were mock-treated (DMSO) or treated with 5 μM AZD6738 and 1 μM NU6140 individually or in combination for 1 h. Points and error bars are the mean and standard error, respectively, of 3 independent experiments.

**Figure S5.**
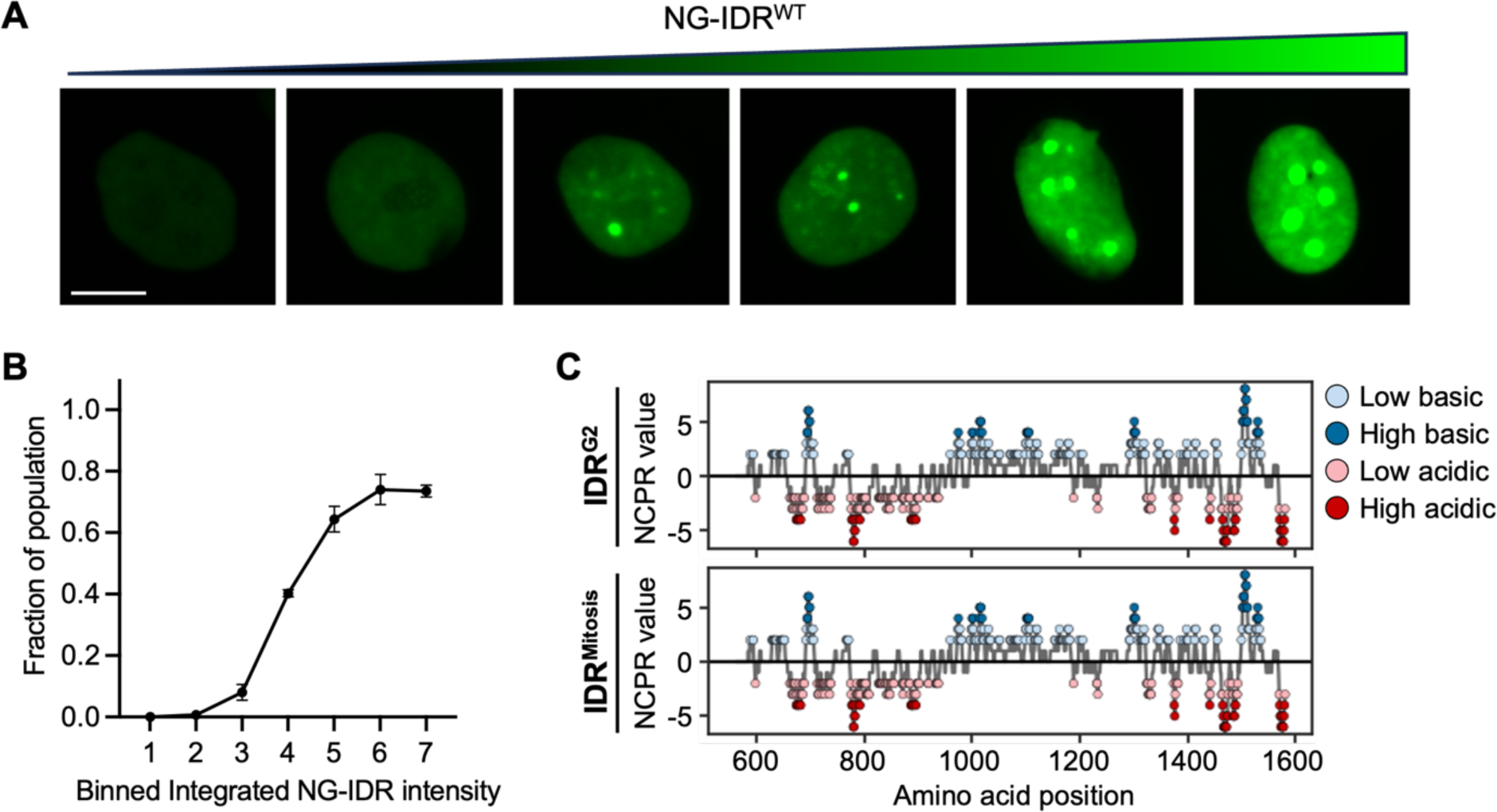
Concentration-dependent formation of mNeonGreen-MED1^IDR^ condensates. (A) Representative images of mNeonGreen-MED1^IDR^ fluorescence in MCF10 cells showing focus formation. Scale bar 10 μm. (B) Line plot showing a threshold expression level when mNeonGreen-MED1IDR foci form. Mean and standard error are shown for 3 independent experiments. (C) Line plots of the Net Charge Per Residue (NCPR) values for the mutated MED1-IDR variants (IDR^G2^ and IDR^Mitosis^). Colored circles indicate the charge status of the bin, light blue for low basic (> 1), dark blue for high basic (> 3), light red for low acidic (< −1), dark red for high acidic (< −3).

**Figure S6.**
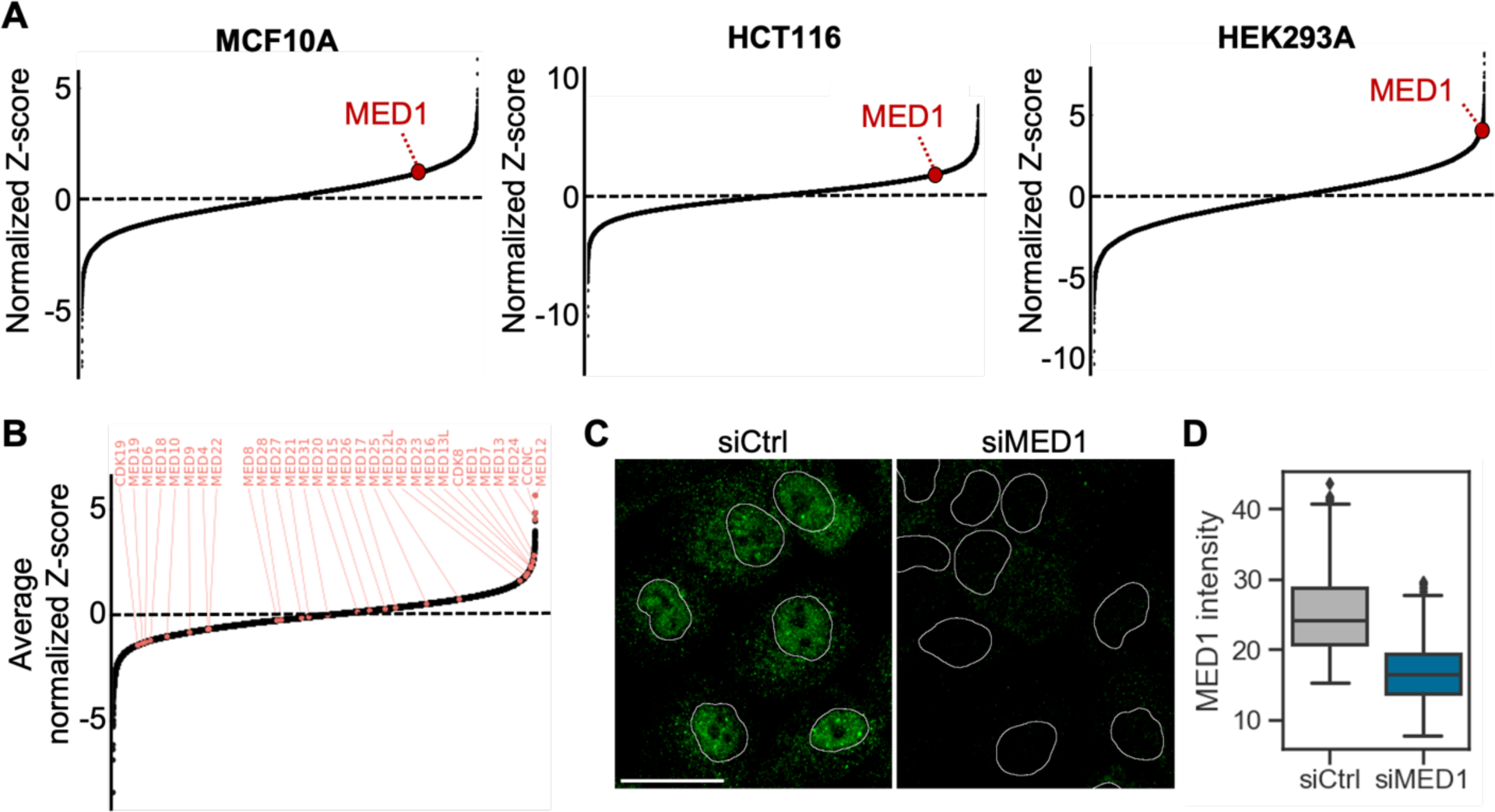
Normalized Z-scores of cell survival with MED1 and other mediator subunits marked in ATR-inhibited cells. (A) Z-score plots showing MED1 in MCF10A, HCT116, and HEK293 cells. (B) Average normalized Z-score in all 3 cell lines from (A) showing the effect of all mediator subunits on ATR-inhibitor response. (C) Representative images of MED1 in MCF10A cells transfected with either a non-targeting siRNA (siCtrl) or siRNA targeting MED1 (siMED1). (D) Box plot showing MED1 intensities in MCF10A cells treated as in (C).

**Figure S7.**
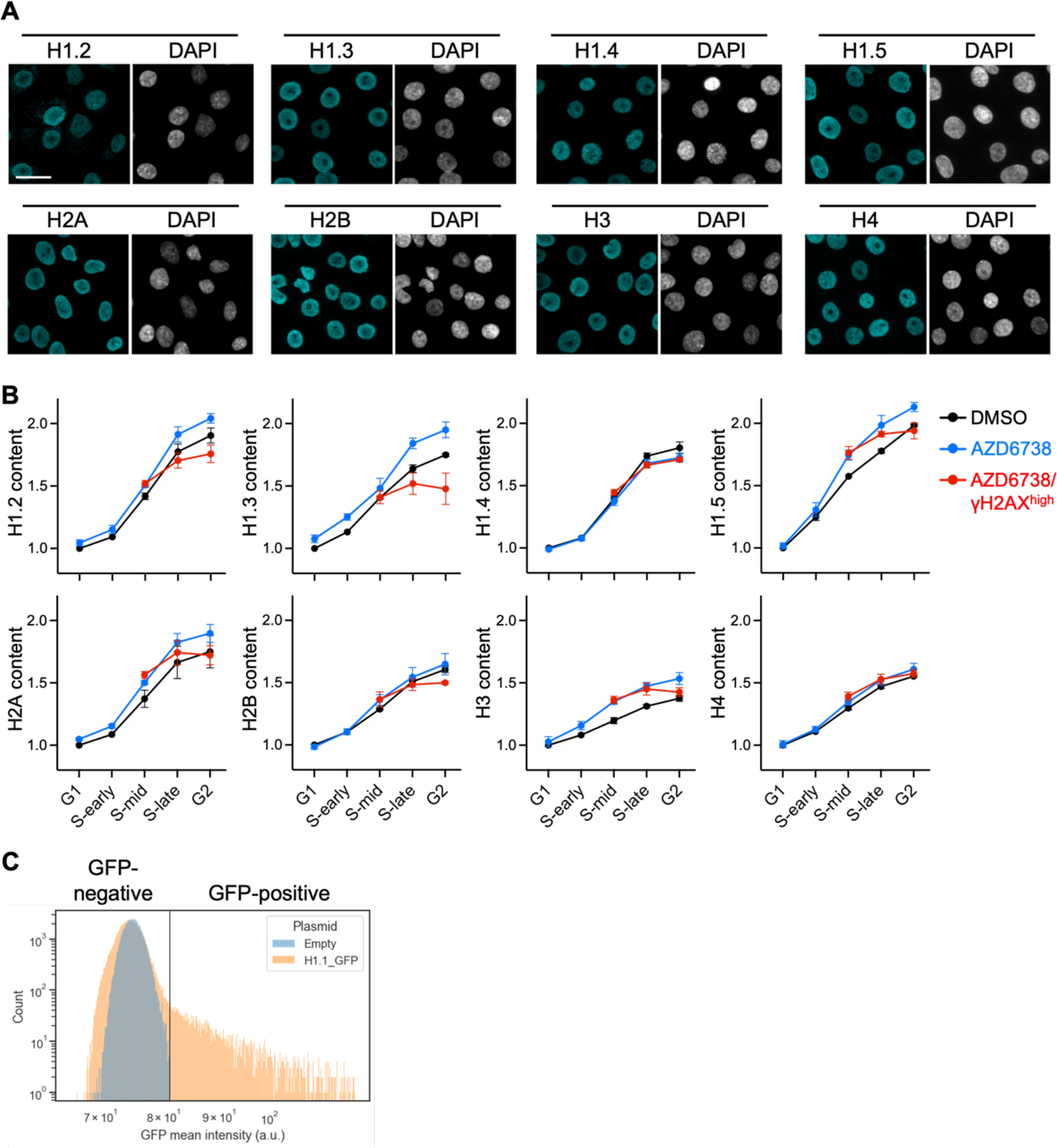
Histone levels across the cell cycle in MCF10A cells. (A) Representative images of the different replication-dependent histones in MCF10A cells. Scale bar 30 μm. (B) Line plots of normalized histone levels across the cell cycle in MCF10A cells. Cells were mock-treated or treated with 5 μM AZD6738 for 16 hours. γH2AX high population was defined as described in Figure 7B. Data show the mean and standard error of 3 independent experiments. (C) Histogram of GFP-negative and GFP-positive cells transfected with H1.1-GFP. Plot is colored to indicate the threshold for defining the negative and positive population.

## REFERENCES

1. Bhat, P., D. Honson, and M. Guttman, Nuclear compartmentalization as a mechanism of quantitative control of gene expression. Nat Rev Mol Cell Biol, 2021. 22(10): p. 653–670.

2. Sabari, B.R., et al., Coactivator condensation at super-enhancers links phase separation and gene control. Science, 2018. 361(6400).

3. Cho, W.K., et al., Mediator and RNA polymerase II clusters associate in transcription-dependent condensates. Science, 2018. 361(6400): p. 412–415.

4. Hnisz, D., et al., A Phase Separation Model for Transcriptional Control. Cell, 2017. 169(1): p. 13–23.

5. Guo, Y.E., et al., Pol II phosphorylation regulates a switch between transcriptional and splicing condensates. Nature, 2019. 572(7770): p. 543–548.

6. Sabari, B.R., A. Dall’Agnese, and R.A. Young, Biomolecular Condensates in the Nucleus. Trends Biochem Sci, 2020. 45(11): p. 961–977.

7. Lyons, H., et al., Functional partitioning of transcriptional regulators by patterned charge blocks. Cell, 2023. 186(2): p. 327–345 e28.

8. Yamazaki, H., et al., Cell cycle-specific phase separation regulated by protein charge blockiness. Nat Cell Biol, 2022. 24(5): p. 625–632.

9. Rai, A.K., et al., Kinase-controlled phase transition of membraneless organelles in mitosis. Nature, 2018. 559(7713): p. 211–216.

10. Harlen, K.M. and L.S. Churchman, The code and beyond: transcription regulation by the RNA polymerase II carboxy-terminal domain. Nat Rev Mol Cell Biol, 2017. 18(4): p. 263–273.

11. Hamperl, S. and K.A. Cimprich, Conflict Resolution in the Genome: How Transcription and Replication Make It Work. Cell, 2016. 167(6): p. 1455–1467.

12. Saldivar, J.C., D. Cortez, and K.A. Cimprich, The essential kinase ATR: ensuring faithful duplication of a challenging genome. Nat Rev Mol Cell Biol, 2017. 18(10): p. 622–636.

13. Saldivar, J.C., et al., An intrinsic S/G(2) checkpoint enforced by ATR. Science, 2018. 361(6404): p. 806–810.

14. Fragkos, M., et al., DNA replication origin activation in space and time. Nat Rev Mol Cell Biol, 2015. 16(6): p. 360–74.

15. Shechter, D., V. Costanzo, and J. Gautier, ATR and ATM regulate the timing of DNA replication origin firing. Nat Cell Biol, 2004. 6(7): p. 648–55.

16. Katsuno, Y., et al., Cyclin A-Cdk1 regulates the origin firing program in mammalian cells. Proc Natl Acad Sci U S A, 2009. 106(9): p. 3184–9.

17. Daigh, L.H., et al., Stochastic Endogenous Replication Stress Causes ATR-Triggered Fluctuations in CDK2 Activity that Dynamically Adjust Global DNA Synthesis Rates. Cell Syst, 2018. 7(1): p. 17–27 e3.

18. Boija, A., et al., Transcription Factors Activate Genes through the Phase-Separation Capacity of Their Activation Domains. Cell, 2018. 175(7): p. 1842–1855 e16.

19. Klein, I.A., et al., Partitioning of cancer therapeutics in nuclear condensates. Science, 2020. 368(6497): p. 1386–1392.

20. Jaeger, M.G., et al., Selective Mediator dependence of cell-type-specifying transcription. Nat Genet, 2020. 52(7): p. 719–727.

21. Toledo, L.I., et al., ATR prohibits replication catastrophe by preventing global exhaustion of RPA. Cell, 2013. 155(5): p. 1088–103.

22. Nizami, Z., S. Deryusheva, and J.G. Gall, The Cajal body and histone locus body. Cold Spring Harb Perspect Biol, 2010. 2(7): p. a000653.

23. Duronio, R.J. and W.F. Marzluff, Coordinating cell cycle-regulated histone gene expression through assembly and function of the Histone Locus Body. RNA Biol, 2017. 14(6): p. 726–738.

24. Tatomer, D.C., et al., Concentrating pre-mRNA processing factors in the histone locus body facilitates efficient histone mRNA biogenesis. J Cell Biol, 2016. 213(5): p. 557–70.

25. Marzluff, W.F., E.J. Wagner, and R.J. Duronio, Metabolism and regulation of canonical histone mRNAs: life without a poly(A) tail. Nat Rev Genet, 2008. 9(11): p. 843–54.

26. Liu, X., et al., Time-dependent effect of 1,6-hexanediol on biomolecular condensates and 3D chromatin organization. Genome Biol, 2021. 22(1): p. 230.

27. Lemmens, B., et al., DNA Replication Determines Timing of Mitosis by Restricting CDK1 and PLK1 Activation. Mol Cell, 2018. 71(1): p. 117–128 e3.

28. Smits, V.A., P.M. Reaper, and S.P. Jackson, Rapid PIKK-dependent release of Chk1 from chromatin promotes the DNA-damage checkpoint response. Curr Biol, 2006. 16(2): p. 150–9.

29. Branigan, T.B., et al., MMB-FOXM1-driven premature mitosis is required for CHK1 inhibitor sensitivity. Cell Rep, 2021. 34(9): p. 108808.

30. Zonderland, G., et al., The TRESLIN-MTBP complex couples completion of DNA replication with S/G2 transition. Mol Cell, 2022. 82(18): p. 3350–3365 e7.

31. Sharma, K., et al., Ultradeep human phosphoproteome reveals a distinct regulatory nature of Tyr and Ser/Thr-based signaling. Cell Rep, 2014. 8(5): p. 1583–94.

32. Olsen, J.V., et al., Quantitative phosphoproteomics reveals widespread full phosphorylation site occupancy during mitosis. Sci Signal, 2010. 3(104): p. ra3.

33. Newcombe, E.A., et al., How phosphorylation impacts intrinsically disordered proteins and their function. Essays Biochem, 2022. 66(7): p. 901–913.

34. Brown, E.J. and D. Baltimore, ATR disruption leads to chromosomal fragmentation and early embryonic lethality. Genes Dev, 2000. 14(4): p. 397–402.

35. Cortez, D., et al., ATR and ATRIP: partners in checkpoint signaling. Science, 2001. 294(5547): p. 1713–6.

36. de Klein, A., et al., Targeted disruption of the cell-cycle checkpoint gene ATR leads to early embryonic lethality in mice. Curr Biol, 2000. 10(8): p. 479–82.

37. Wang, C., et al., Genome-wide CRISPR screens reveal synthetic lethality of RNASEH2 deficiency and ATR inhibition. Oncogene, 2019. 38(14): p. 2451–2463.

38. Buisson, R., et al., Distinct but Concerted Roles of ATR, DNA-PK, and Chk1 in Countering Replication Stress during S Phase. Mol Cell, 2015. 59(6): p. 1011–24.

39. Marzluff, W.F., et al., The human and mouse replication-dependent histone genes. Genomics, 2002. 80(5): p. 487–98.

40. Spegg, V., et al., Phase separation properties of RPA combine high-affinity ssDNA binding with dynamic condensate functions at telomeres. Nat Struct Mol Biol, 2023. 30(4): p. 451–462.

41. Du, M., et al., Direct observation of a condensate effect on super-enhancer controlled gene bursting. Cell, 2024. 187(2): p. 331–344 e17.

42. Singh, R.K., et al., Excess histone levels mediate cytotoxicity via multiple mechanisms. Cell Cycle, 2010. 9(20): p. 4236–44.

43. Willcockson, M.A., et al., H1 histones control the epigenetic landscape by local chromatin compaction. Nature, 2021. 589(7841): p. 293–298.

44. Thorslund, T., et al., Histone H1 couples initiation and amplification of ubiquitin signalling after DNA damage. Nature, 2015. 527(7578): p. 389–93.

45. Prendergast, L. and D. Reinberg, The missing linker: emerging trends for H1 variant-specific functions. Genes Dev, 2021. 35(1-2): p. 40–58.

46. Uggenti, C., et al., cGAS-mediated induction of type I interferon due to inborn errors of histone pre-mRNA processing. Nat Genet, 2020. 52(12): p. 1364–1372.

47. Zou, L. and S.J. Elledge, Sensing DNA damage through ATRIP recognition of RPA-ssDNA complexes. Science, 2003. 300(5625): p. 1542–8.

48. Egger, T., et al., Spatial organization and functions of Chk1 activation by TopBP1 biomolecular condensates. Cell Rep, 2024. 43(4): p. 114064.

49. Frattini, C., et al., TopBP1 assembles nuclear condensates to switch on ATR signaling. Mol Cell, 2021. 81(6): p. 1231–1245 e8.

50. Lalonde, M., et al., An automated image analysis pipeline to quantify the coordination and overlap of transcription and replication activity in mammalian genomes. Methods Cell Biol, 2024. 182: p. 199–219.

